# Characterization of *FLOWERING LOCUS T* related genes and their putative gene regulatory network in semi-winter *Brassica napus* cultivar Zhongshaung11

**DOI:** 10.1101/2024.11.13.623477

**Authors:** Juanjuan Wang, Hao-Ran Zhou, Petra Tänzler, Na Ding, Jing Wang, Franziska Turck

**Affiliations:** Max Planck Institute for Plant Breeding Research, Carl von Linné Weg 10, 50829 Köln; MOA Key Laboratory of Crop Ecophysiology and Farming System in the Middle Reaches of the Yangtze River, College of Plant Science and Technology, Huazhong Agricultural University, Wuhan, PR China

**Keywords:** FLOWERING LOCUS T, *Brassica napus*, flowering time, transcriptional regulation, CONSTANS, NF-Y

## Abstract

In many species, *FLOWERING LOCUS T* (*FT*)-like genes promote the floral transition by integrating environmental signals, in particular photoperiod, and internal cues. Here we show that *Brassica napus* contains 6 *FT*-like genes and 2 pseudogenes belonging to 3 orthogroups. All *B. napus FT*-like genes induce early flowering when expressed at the shoot apical meristems of *Arabidopsis thaliana ft* mutants, but *BnaFT.C6* and non-orthologous *FT*-like genes do not encode fully functional mobile florigens. In the case of *BnFT.C6*, the functional change is associated with a T to C amino acid change that is restricted to semi-winter accessions. Expression of orthologs of *FT* is photoperiod-dependent and two distal enhancers are conserved; however, the homeologs *BnaFT.A7* and *BnaFT.C6* show rearrangements of DNA motifs binding NF-Y/CO and NF-Y transcriptional activator complexes between the promoter and downstream enhancers. Motif rearrangements correlate with differences in tissue-specific expression. Furthermore, homeologs with rearranged motifs could not be trans-activated by *B. napus* CO in transient assays although they genes show LD photoperiod-dependent expression. We propose that differential diurnal expression of *NF-Y* genes contributes to the photoperiod-dependent regulation of *B. napus FT* genes.

**Significance statement:** *B. napus FT*-like genes are a case study for gene retention and retraction after multiple rounds of polyploidisation. Retained *B. napus FT*-like genes show preferred expression in different developmental stages and amino-acid changes that affect florigen function.

## Introduction

Flowering is an important growth stage in plants with direct impact on biomass, survival and seed production and hence crucial for crop yield. Flowering is promoted by environmental and developmental signals in angiosperms and based on observations made in the model plant *Arabidopsis thaliana,* the underlying flowering promoting pathways have been categorised as the vernalization, photoperiod, autonomous, gibberellin (GA) and age pathways (Andrés and Coupland, 2012; Teotia and Tang, 2015). A major integrator of flowering cues in *A. thaliana* is *FLOWERING LOCUS T* (*FT*), which plays a pivotal role in the photoperiod pathway but also senses cues from other pathways (Bratzel and Turck, 2015; Takagi *et al*., 2023). FT acts as a mobile florigen, as it is transcribed in the phloem companion cells of leaves in response to a long day (LD) photoperiod. This subsequently allows FT protein to move as a protein through sieve elements to the shoot apical meristem (SAM), where it associates with the transcription factor FLOWERING LOCUS D (FD). The FT/FD complex directly promotes the expression of genes that switch the developmental program of the SAM from vegetative to reproductive growth (Abe *et al*., 2005; Wigge *et al*., 2005; Jaeger and Wigge, 2007).

*FT*-like genes were shown to promote flowering in many plant species; furthermore, as mobile signals they can also impact inflorescence commitment and architecture, shoot branching, storage organ formation and seed dormancy, underscoring their importance for plant fitness (S., Jin *et al*., 2021). Many plant species show *FT*-like gene expansion, such as the *FT* paralog *TWIN SISTER OF FT* (*TSF*) in *A. thaliana*, which has diversified in expression but not protein function (Yamaguchi *et al*., 2005). Functional diversification of paralogs has also been reported; for example, *BvFT2* and *BvFT1* in *Beta vulgaris* (sugar beet) respectively activate and repress flowering, which is crucial to integrate vernalization to flowering regulation in this species (Pin *et al*., 2010).

Research on the transcriptional regulatory mechanisms of *FT* uncovered a network of activating and repressing transcription factors, epigenetic regulators and cis-acting regulatory elements (Takagi *et al*., 2023; Bratzel and Turck, 2015). *FT* activation in LD requires the combined input of the proximal promoter and two distal transcriptional enhancers, located at 5.7-kb and 1.6-kb distance up-and down-stream of the structural gene, respectively (Liu *et al*., 2014; Cao *et al*., 2014; Adrian *et al*., 2010; Tiwari *et al*., 2010; Zicola *et al*., 2019). *FT* regulatory regions recruit related protein complexes: the canonical ternary eukaryotic CCAAT box binding factors (CBF), also known as NF-Y complexes and related plant-specific NF-Y/CO complexes. NF-Y complexes bind CCAAT motifs and are composed of one of each NF-YB, NF-YC and NF-YA subunits.

In NF-Y/CO complexes, the NF-YA component is replaced with the plant-specific CONSTANS, CONSTANS-like, TOC1 (CCT) domain protein CONSTANS (CO). CCT domains exhibit structural similarity to NF-YA, yet demonstrate direct, high-affinity binding to CCACA in lieu of CCAAT motifs (Wenkel *et al*., 2006; Gnesutta *et al*., 2017; Lv *et al*., 2021).

Photoperiod regulation in plants involves transcriptional and post-transcriptional control of CO. Transcriptional control includes the degradation of CO’s repressors, such as CYCLIC DOF FACTOR 1-LIKE (CDFs), by a blue light receptor complex that contains FKF1 and GIGANTEA (GI) (Fornara *et al*., 2009; Imaizumi *et al*., 2005; Sawa *et al*., 2007). CO transcription is repressed in the morning and activated in the late afternoon. CO protein decreases in the night due to CONSTITUTIVE PHOTOMORPHOGENESIS 1 (COP1) and SUPPRESSOR OF PHYA-105 (SPA) 1-4 proteins (Laubinger *et al*., 2006; Jang *et al*., 2008), while photoreceptors regulate CO stability throughout the day (Valverde *et al*., 2004; Yanovsky and Kay, 2002; Posé *et al*., 2013). This regulation allows CO protein to accumulate at dusk under LD but not short day (SD) photoperiod. Additionally, other transcription factors like PHYTOCHROME INTERACTING FACTORs (PIFs) and CRYPTOCHROME 2-INTERACTING bHLH (CIB) proteins modulate *FT* expression in response to light and temperature (Galvāo *et al*., 2019; Fernández *et al*., 2016; Kumar *et al*., 2012; Liu *et al*., 2013; Liu *et al*., 2018).

A strong motivation for in-depth studies in genetic model organisms is the potential for translational application to crops. While key pathway integrators are mostly conserved, how these are embedded in gene regulatory networks can diverge even between closely related species. The Brassica genus includes important crops, which diverged from *A. thaliana* 27 million years ago. The genus underwent a genome triplication and differentiated three lineages, designated as type A, B and C genomes (Kagale *et al*., 2014). Detailed comparison of synteny between and within these genomes revealed the presence of a least fragmented (LF) genome and two genomes that were mildly (MF1) or mostly fragmented (MF2) with rearrangements often resulting in gene loss (Parkin *et al*., 2005; Cheng *et al*., 2013). Hybrids between Brassica lineages were strongly favoured by early farmers in their selection of crops. The most important, *Brassica napus* (AACC, 2n=38) is mostly grown as oilseed rape (canola) and originated around 7,500 years ago through hybridization between the diploid progenitors *Brassica rapa* (AA, 2n=20) and *Brassica oleracea* (CC, 2n=18), probably in the Mediterranean region (Wu *et al*., 2019; Chalhoub, 2014; Lu *et al*., 2019). The global *B. napus* gene pool has undergone several post-domestication ecogeographic radiations, which provided the origin of three cultivars according to their vernalization requirements (Zou *et al*., 2019). Winter rapeseed is mainly found in Europe and requires a prolonged period of low temperatures to transit from vegetative to reproductive growth; semi-winter rapeseed, mainly produced in China, can initiate flowering after a shorter vernalization period; and spring rapeseed, which presents a wide distribution in Northern Europe, Canada, and Australia, can flower and reproduce without vernalization.

Previous studies have shown that gene groups involved in flowering regulation in *A. thaliana* show both expansion and retraction in *B. napus*, indicating that purifying and diversifying selection takes place. In the case of *FT*, six or seven homologs were identified, depending on study and genetic background (Schiessl *et al*., 2014; Schiessl, 2020). Notably, silencing of a *FT* syntenic *B. napus* gene on chromosome C02 (*BnaFT.C2*) coincided with two TE insertions between the proximal promoter and consequent high levels of cytosine methylation in TE sequences within proximal promoter region (J., Wang *et al*., 2012). Three *BnaFT* paralogues, designated as *BnaA2.FT*, *BnaC6.FT.a* and *BnaC6.FT.b* were associated with two major QTL clusters for flowering time, in a mapping population from the cross of the winter-type cultivar Tapidor and the semi-winter cultivar NY7 (Wang *et al*., 2009). Analysis of selective sweeps between spring, semi-winter and winter ecotypes, and GWAS for flowering time confirmed that *BnaFT.A2* (*BnaA02g12130D*) is significantly associated with flowering-time variation (Wu *et al*., 2019). In addition, EMS-induced mutant alleles of *BnaC6.FT.b* exhibited delayed flowering compared to the control group, while mutants of *BnaC6.FT.a* exhibited flowering patterns akin to the non-mutated parent in winter-type inbred line Express 617 (Guo *et al*., 2014). Furthermore, loss-of-function mutants in Westar and RNAi lines in Tapidor of *BnaFT*.A2 have smaller leaves, a lower net photosynthetic rate, as well as a substantial delay in flowering time compared to their parents (Jin *et al*., 2022).

Here we present a comparative analysis of six *FT*-like genes encoded by the semi-winter *B. napus* cultivar Zhongshaung11 (ZS11).

## Materials and Methods

### Plant material and growth conditions

*A. thaliana* accession Columbia (Col-0) and the *ft-10* T-DNA insertion mutant (GABI-Kat line 290E08) were cultivated in a temperature-controlled greenhouse in LD photoperiod (20-24°C, 16h light/8h dark) on soil. *B. napus* cultivar ZS11 plants were grown in the greenhouse (20-24°C) for the collection of tissues from different growth stages and in growth chambers (CLF Plant Climatics, AR-95L3X, 60% light intensity (12570 LUX), 75% relative humidity. 22°C) under LD (16 h light /8 h dark) or SD (8 h light /16 h dark) for the collection of leaf material for diurnal transcriptomes. For vernalization, plants were moved to vernalization rooms (4°C, SD).

### Identification of *BnaFT* and *NF-Y* homologs in *B. napus*

Synteny analysis was performed using the MCscan algorithm (Y., Wang *et al*., 2012) as implemented in the JCVI toolkit (Tang *et al*., 2024). Genome sequences and genome annotations for *B. napus* (ZS11.v0) and *Brassica oleracea* var capitata (JZS.v2) (BnPIR; http://cbi.hzau.edu.cn/bnapus) <u>(Song *et al*., 2020)</u>, *Brassica rapa*Z1_v2 (https://www.genoscope.cns.fr/externe/plants/data/), *Schrenkiella parvula* v2.2 (Phytozome13; https://phytozome-next.jgi.doe.gov/) and *A. thaliana* Col-0 Araport11 (TAIR; https://www.arabidopsis.org/) were downloaded from the indicated sources. Amino acid sequences of annotated proteins were generated using the scripts of the JCVI toolkit and blasted all against all using the LAST algorithm (Kiełbasa *et al*., 2011). To identify syntenic genes, 6 iterations of comparing all genomes against the *A. thaliana* and *S. parvula* genomes by MCscan were performed.

Homologs of FT (AT1G65480) (Supplemental Table 1) and NF-Y proteins (Supplemental Table 2) in *B. napus* var ZS11 were identified using screen criteria of E-value ≤ 1.0E-20, coverage ≥ 95%, identity ≥ 80%. *FT*-like pseudogenes were identified by relaxing the coverage criteria. To include them in the alignment, frameshift mutations and stop codons were corrected according to the output of the Scipio webtool (Hatje *et al*., 2011). To construct a phylogenetic tree of FT-like proteins, full length protein sequences of FT, TSF, TFL1, BFT and MFT from *A. thaliana* were obtained from TAIR database (https://www.arabidopsis.org/). Protein alignments were performed with ClustalW and a 60% majority-rule consensus tree was constructed from a bootstrapped (n=1000) neighbor joining (NJ) using MEGA11 (Tamura *et al*., 2021). NF-Y proteins were aligned and visually curated with ClustalW. An maximum-likelihood phylogenetic tree was generated that included *A.thaliana* and *B.napus* homologs of all three NF-Y families for overview.

### mVISTA analysis

Genomic sequences including upstream and downstream sequences up to the next flanking genes were submitted to the mVISTA webtool for alignment (https://genome.lbl.gov/vista/mvista/submit.shtml).

### Annotation of cis-regulatory motifs at conserved regulatory regions

Sequences identified as aligning to conserved Blocks C, A and E were extracted and re-aligned using ClustalW in MEGA11 (Tamura *et al*., 2021). Motifs were annotated and plotted using custom scripts in R.

### Plasmids construction for complementation and transient expression

The vector *(C+A)-FTpromoter(FTp)-FTcDNA-pGreen* and transgenic *(C+A)-FTp-FTcDNA-pGreen*/*ft-10* plants were described previously (Liu *et al*., 2014). Full length coding sequences of *BnaFT.A2*, *BnaFT.C2*, *BnaFT.A7*, *BnaFT.C6*, *BnaNFT.A7*, and *BnaCFT.C4* were amplified from ZS11 cDNA using primer pairs as indicated in Supplemental Table 3. PCR products were introduced into *(C+A)-FTp-FTcDNA-pGreen* (Liu *et al*., 2014) via *Hind*III and *Sac*I restriction enzyme sites to replace the original *FT* CDS on the backbone using NEBuilder® HiFi DNA Assembly Master Mix according manufacturer’s instructions (NEB, Frankfurt, Germany). Similarly, the mutant versions of *BnaFT.A7* and *BnaFT.C6*, namely *BnaFT.A7*m and *BnaFT.C6*m, were amplified using mutagenic primers and *(C+A)-FTp-BnaFT.A7-pGreen* and *(C+A)-FTp-BnaFT.C6-pGreen* as the templates.

For expression of *FT*-like genes at the shoot apex, the vector *FDp-FD-3xVENUS-pER8* was used as backbone. Full-length coding sequences were amplified with two rounds of PCR to add an N-terminal HA-epitope to the full-length sequence using primers as indicated in Supplemental Table 3. The final PCR products were inserted into vector backbone FDprom-FD3V-pER8 using *Xho*I restriction enzyme sites using NEBuilder® HiFi DNA Assembly Master Mix.

For green luciferase (LUC68) reporter vectors, promoters of *BnaFT.A2*, *BnaFT.A7*, *BnaFT.C6* and two fragments of *BnaFT.C2* corresponding to Block C and Block A were amplified from ZS11 gDNA using primer pairs listed in Supplementary Table 3 and introduced via *Sca*I and *Nco*I restriction enzyme sites to replace the *Block A FT* promoter in *BlockA-FTp-LUC68-pGreen* using the NEBuilder® HiFi DNA Assembly Master Mix. *BlockA-FTp-LUC68-pGreen, 5.7Kb-FTp-LUC68-pGreen* vectors were described previously (Adrian *et al*., 2010). For effector vectors, the cassettes *35Sp-BnaCO.A10, 35Sp-BnaCO.C9*, *35Sp-CO* and *35Sp-H2B* were introduced into the vector *BlockA-FTp-LUC68-pGreen* via restriction enzyme sites *Sca*I and *Sac*I using the NEBuilder® HiFi DNA Assembly Master Mix, to replace *BlockA-FTp-LUC68*. The vector *35Sp-RedLUC-pJAN33* and *35Sp-LUC68-pJAN33* for normalization and filter correction were previously described (Adrian *et al*., 2010).

### Plant transformation

Plasmids were introduced into *Agrobacterium tumefaciens* strain GV3101(pSoup) to transform *ft-10* by floral dipping (Clough and Bent, 1998). Transformants were selected on soil by applying Basta (glufosinate) spray (0.1%) twice for the vectors using *(C+A)-FTp-FTcDNA-pGreen* and on plates supplemented with hygromycin (50 mg/L) for the vectors using *FDp-FD3V-pER8* as the backbone.

#### Tobacco infiltration and luciferase assay

Suspensions of *A. tumefaciens* GV3101(pSoup) strain carrying the plasmids of interest were mixed according to the experimental design and then infiltrated into the underside of *N. benthamiana* leaves as described (Sparkes *et al*., 2006) with the following adjustments: 10 mM MES and 40 µM acetosyringone were added to the *A. tumefaciens* culture medium, which was then incubated overnight at 28℃. The culture was centrifuged at 4,000 *g* for 15 min and the supernatant was replaced with the infiltration medium to adjust the optical density at 600nm to 2.0. Cultures were then placed at room temperature for at least 1 h before being used for infiltration. Infiltrated plants were cultivated overnight in a dark humidified cabinet and then transferred to the greenhouse for 48h. Detection of green luminescent LUC68 and red luminescent RedLUC *in vivo* was performed as described (Adrian *et al*., 2010). Briefly, leaves were excised, placed on moist filter paper, sprayed with 1mM luciferin dissolved in 0.25% Tween20 and incubated for 1min. Using the digital camera of a LAS4000 device (GE Healthcare), LUC68 and RedLUC signals were integrated sequentially for 2 min through 560nm short pass and 610nm long pass camera lens filters, respectively. To calculate the amount of RedLUC activity measured through the green filter and *vice versa*, bombardments with only *35Sp-RedLUC-pJAN33* or *35Sp-LUC68-pJAN33* were performed. The ratios of signals were calculated using a custom script in R based on the Chroma-Luc^TM^ calculator provided by Promega. Bioluminescence signals were quantified from 16bit greyscale images in TIFF format using ImageLab software (Biorad).

### RNA extraction, reverse transcription and quantitative reverse transcribed-PCR (qRT-PCR)

Total RNA from *B. napus* and *A. thaliana* plants was extracted using the RNeasy Mini Kit (Qiagen, Cat. no. 74104) following manufacturer’s instructions. For each sample, 4 µg of total RNA was treated with DNA-free^TM^ Kit (Invitrogen, REF AM1906), and cDNA synthesis was performed with Superscript^TM^ IV reverse transcriptase kit (Invitrogen, REF18090050) following manufacturer’s instructions. qRT-PCR was conducted in three technical replicates for each biological replicate using the Bio-Rad CFX384^TM^ system (Bio-Rad), technical replicates were collapsed to their median prior statistical analysis.

For the analysis of tissue-specific expression, *B. napus* cultivar ZS11 were cultivated for 4 weeks at 22°C in the greenhouse under LD condition, vernalization for 4 weeks at 4°C in SD, and transferred back to the LD greenhouse. Throughout the developmental progression, sequential samplings of three biological replicates occurred at Zeitgeber time 12 (ZT12). Root and leaf 6 were sampled prior to vernalization, leaf 8, paraclade leaf, floral bud, flower and siliques were sampled after vernalization. Material from leaf 8 was also sampled from non-vernalized plants.

For diurnal expression analysis, ZS11 was cultivated for four weeks in separate LD or SD growth chambers, followed by an intervening vernalization period of 4 weeks, after which plants were transferred back to the LD or SD chambers, respectively. Leaf 6 was sampled just before plants were transferred to the vernalization and leaf 8 was harvested one week after transfer to LD or SD chambers at 22°C. Biological triplicates were sampled every 4 hours for 24 hours. The expression levels of *BnaFT.A2*, *BnaFT.C2*, *BnaFT.A7*, *BnaFT.C6*, *BnaNTF.A7*, *BnaCTF.C4*, *BnaCO.A10*, *BnaCO.C9* and *BnaENTH* in *B. napus* were tested with gene-specific primer pairs that are listed in Supplemental Table 3. Standard curves of each pair of primers were obtained through PCR using gradient-diluted ZS11 genome DNA as the template and Ct values were adjusted using amplification efficiencies prior to using the-ddCt method. The *BnaENTH* gene was used to correct for differences in cDNA concentration, a dilution of genomic DNA was used as common scale for all genes.

To measure expression of complementation lines, leaf material was harvested at ZT16 from 14-day old *A. thaliana* plants grown in soil in LD at 22°C in greenhouse conditions. To quantify expression of transgenes in *A. thaliana*, specific primers pairs against *FT*, *BnaNFT.A7*, *BnaCFT.C4* and a common pair for four *BnaFT* genes were titrated against the plasmids used for transformation. A primer pair specific to the phosphinothricin resistance marker (*BAR*) was used to normalize plasmid concentrations, primers against *PROTEIN PHOSPHATASE 2A* (*PP2A*) were used to correct for differences in cDNA amount (see Supplemental Table 3). Expression levels were related to the median *FT* expression in Col-0.

### Generation and analysis of RNAseq data

Total RNA from three biological replicates of leaf 8 from 24 h time-courses of vernalized *B. napus* var ZS11 plants grown in LD and SD were sent to BGI (Hongkong, China) for library preparation and sequencing using the DNBSEQ platform. Reads were aligned to the reference transcriptome using STAR version 2.7.10b (Dobin *et al*., 2013) and gene expression was quantified as transcript per million (TPM) using RSEM version 1.3.3 (Li and Dewey, 2011). Diurnally expressed genes were identified using Bioconductor R libraries DESeq2 (Dobin *et al*., 2013) and RAIN (Thaben and Westermark, 2014).

## Results

### The *B. napus* genome carries four functional *FT* orthologous and two *FT* paralogous genes

A protein similarity search against a high-quality assembly of *B. napus* semi-winter cultivar ZS11 (Song *et al*., 2020) detected eight FT homologs with identity scores above 81%, of which two were annotated as homologs of TSF but covered the query sequence only partially (Supplemental Table 1). A query of the genomic sequence with the Scipio webtool (Hatje *et al*., 2011) confirmed the presence of six complete *FT-*like genes while proteins showing an incomplete overlap are likely pseudo-genes due to presence of frame shifts and/or non-canonical introns in the predicted coding sequence (Supplemental Table 3, Supplemental Figure 1).

We compared the linearity of lastal hits between *A. thaliana, S. paruvla*, and *B. napus* genomes using the MCscan toolkit (Tang *et al*., 2008). *S. parvula* is a diploid representative of the Brassicacea lineage II, which includes the genus Brassica. The analysis confirmed that four *FT* syntenic genes are located on chromosomes A02, C02, A07, and C06 (Wang *et al*., 2009) (Figure 1A, Supplemental Data 1). A *TSF* ortholog was present in *S. parvula* but missing at the corresponding syntenic blocks in *B. napus*, indicating gene loss in the genus Brassica rather than a lineage I specific duplication (Figure 1B). On the other hand, *S. parvula* and *B. napus* featured an *FT-*like gene at a novel syntenic position on chromosome A07, for which no correspondence could be found in *A. thaliana* (Figure 1C). This gene is a homeolog of both *FT*-like pseudogenes. We propose the name *NEW SISTERS OF FT AND TSF* (*NFT*) to distinguish them from *FT* or *TSF* orthologous genes. *BnaFT.A7*/*BnaNFT.A7* and *BnaFT.C6*/Pseudo-*BnaNFT.C6* are located near a region showing a large inverted duplication in Chromosomes A07/C06 (Parkin *et al*., 2005); however juxtaposition with the *FT* and *NFT* syntenic segments showed that the *FT* gene is situated at the margin of the original syntenic fragment, narrowly avoiding the duplication (Supplemental Figure 2). A sixth functional *FT*-like gene was located on chromosome C04 at a position unique to *B. napus* and the C-genome parent *B. oleracea*, but absent from *B. rapa*, *S. parvula* and *A. thaliana* (Figure 1D). We propose the name *C-GENOME SISTER OF FT AND TSF* (*CFT*) for this gene.

**Figure 1.**
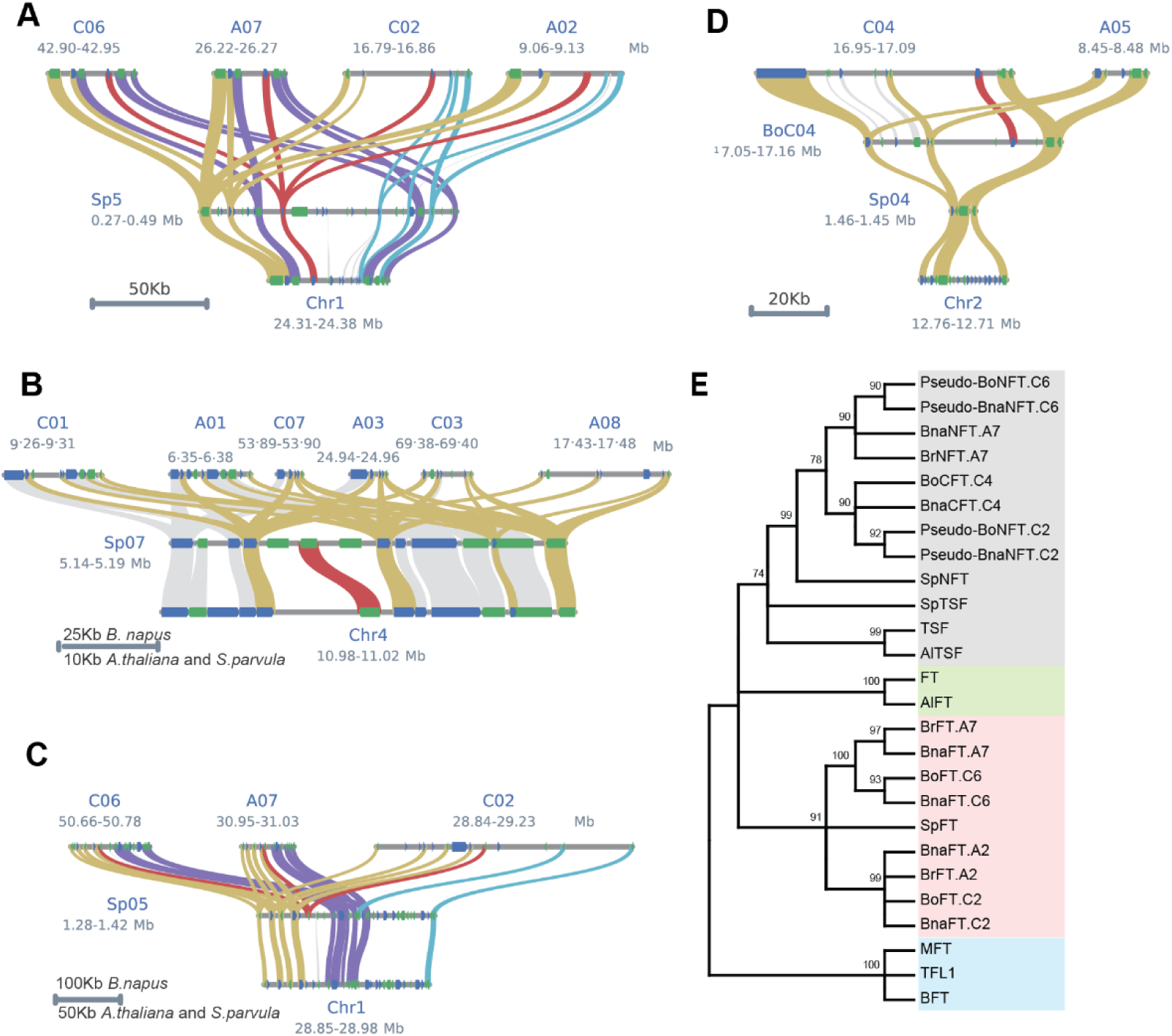
Phylogenetic and syntenic analysis of *FT* and *TSF* and their homologues in *B. napus*, *B*. *oleracea* and *S. parvula*. (A)-(D) Microsynteny between *FT* homologs*. FT-*like genes from *A. thaliana* (bottom), *S. parvula* (middle) and *B. napus* (top) are connected by red lines, orthologs of flanking genes present in all genomes are connected by golden lines, orthologs of flanking genes that exist only in a subset of Brassica genomes are connected by purple, blue and gray lines. Genes are depicted as boxes (blue forward, green reverse orientation). **(A)** Four paralogous *FT* orthologs were identified in *B. napus*, one in *S. parvula*. **(B)** One *TSF* ortholog was identified in *S. parvula*, none in *B. napus*. **(C)** Three paralogous *FT-* like genes (*BnaNFT*) on Chr. A07/C02/C06 in *B. napus* have an ortholog in *S. parvula*, but not in *A. thaliana*. Only the paralogue on chromosome A07 (*BnaNFT.A7*) encodes a functional gene. **(D)** An *FT-*like gene on Chr. C04 (*BnaCFT.C4*) in *B. napus*, is present in the C-genome parent *B. oleracea* and absent in the *S. parvula* and *A. thaliana* genomes. (**E)** Full-length protein sequences of FT, TSF and FT homologues obtained by blast analysis against *B. napus*, *B. rapa*, *B. oleracea, A. lyrata* and *S. parvula* genomes were used to create the neighbour-joining (NJ) consensus tree using MEGA-11 with 1,000 bootstraps. TFL1, BFT, MFT were set as an outgroup, and FT and TSF orthologues in *A. thaliana* and *A. lyrata* were added for reference.

A bootstrapped neighbour-joining phylogenetic tree was constructed from the alignment of six functional FT-like proteins, two frame-corrected pseudogenes and FT homologs from *B. rapa*, *B. oleracea*, *S. parvula, A. lyrata* and *A.thaliana*. TERMINAL FLOWER 1 (TFL1), BROTHER OF FT AND TFL1 (BFT) and MOTHER OF FT AND TFL1 (MFT) were added as outgroups (Figure 1E). In this tree, TSF/*Al*TSF clustered together with TSF from *S. parvula* and NFT/CFT proteins indicating a more recent duplication history for non-*FT* orthologs (Figure 1A).

### Six extant *B. napus FT* homologs show divergence in their potential to act as mobile florigen

To elucidate the potential of FT-like proteins from *B. napus* to act as mobile florigen, the corresponding cDNAs were stably transformed into the *A. thaliana ft-10* mutant under the control of a modified *FT* promoter (*(C+A)-FTp*) that was previously shown to drive transgene expression in phloem companion cells similar to endogenous *FT* in LD photoperiod (Figure2A) (Liu *et al*., 2014). First, flowering time of at least 4 independent segregating T2 lines was compared by comparing the median values of total leaves (Supplemental Figure 3A and B). These results indicated two distinct complementation groups with *BnaFT.A2*, *BnaFT.A7* and *BnaFT.C2* complementing *ft-10* with delay of 5 to 10 leaves compared to *FT* control lines while *BnaNFT.A7,BnaCFT.C4* and *BnaFT.C6* transgenic lines flowered almost as late as *ft-10*. In order to exclude that expression differences were the cause of the observed variation in complementation, we quantified transgene expression in T3 lines with single locus insertions (Supplemental Figure 3C and D). Signals were quantified against the plasmids used for transformation to allow the direct comparison of expression levels between genes. Although large variation in transgene expression explained outliers within genotypes, lines within +/-4-fold expression band compared to *FT* in Col-0 showed no correlation between expression and flowering time (Supplemental Figure 3C and D). The analysis of two representative T2 lines per genotype confirmed the presence of four complementation groups ranging from *FT* showing full complementation, followed by *BnaFT.A2/BnFT.A7/BnaFT.C2*, then *BnaFT.C6/BnaNFT.A7/BnaFT.C4* with increasingly partial complementation and, last, *ft-10* (Figure 2B). In contrast, when *B. napus FT-*like genes were expressed directly in the SAM under the control of the *FD* promoter (*FDp*) (Figure 2A)(Abe *et al*., 2005), independent T1 lines expressing *FT* or *B. napus FT* homologs flowered earlier than wild type controls, although *FDp-BnaCFT.C4* lines flowered slightly later than all other lines (Figure 2C).

**Figure 2.**
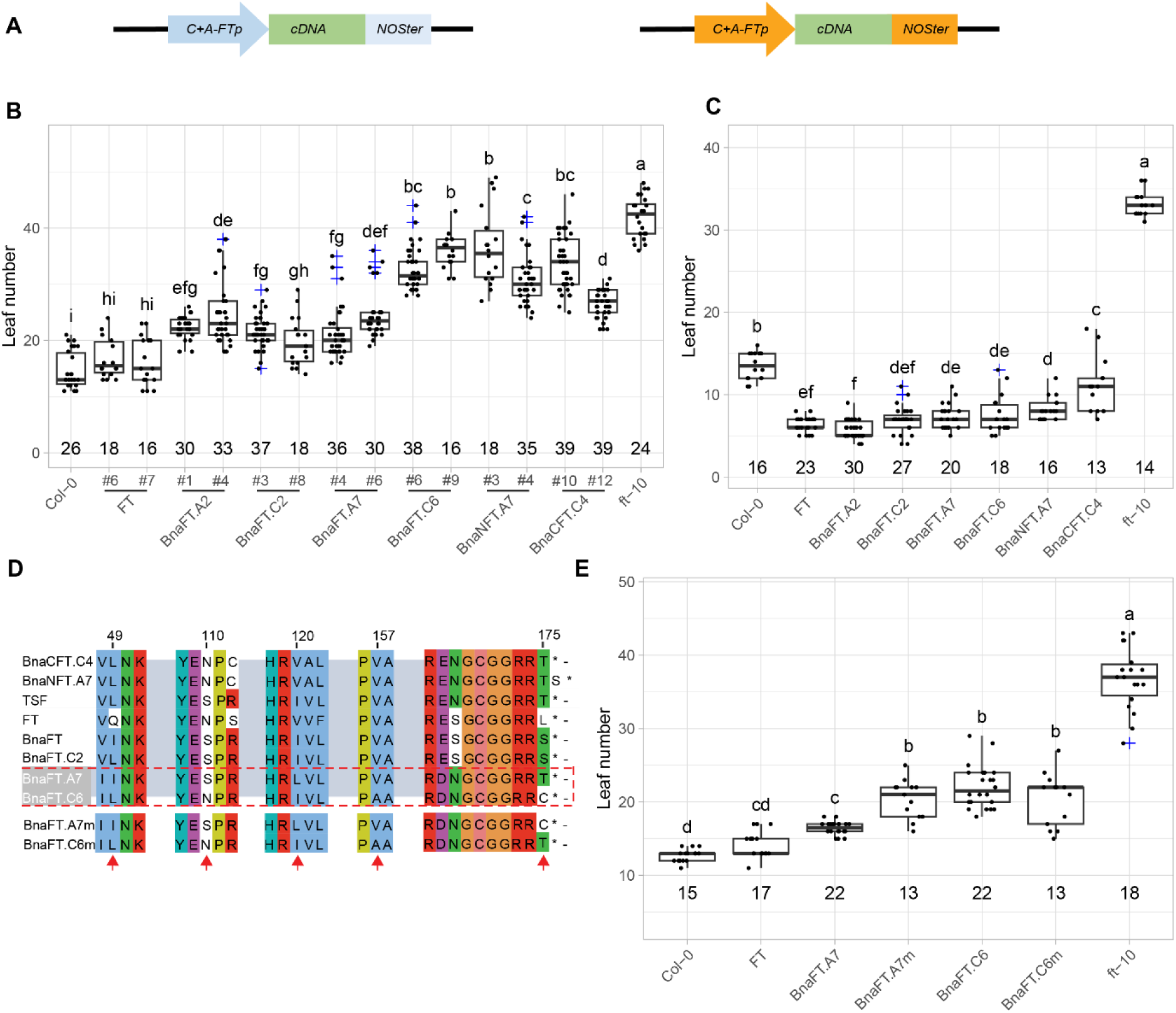
Florigen function of *B. napus FT* homologs. **(A)** Scheme of expression cassettes used for complementation test. **(B)** Flowering time as total leaf number of T3 plants of two representative *ft-10* lines transformed with *FT* and *B. napus FT* homologs driven by *(C+A)-FTp*. The number of plants used for analysis is indicated above the x-axis. **(C)** Flowering time of independent T1 *ft-10* plants transformed with *FT* and *B. napus FT* homologs under the control of *FD*p. The number of independent T1 plants used for analysis is indicated above the x-axis. **(D)** Selection of amino-acids alignment of *A. thaliana* and *B. napus FT* homologs. Arrows indicate differences between *BnaFT.A7* and *BnaFT.C6*. *BnaFT.A7m* and *BnaFT.C6m* are mutant versions in which the last amino acid residue was swapped. **(E)** Flowering time of independent T1 *ft-10* T1 plants transformed with *BnaFT.C6, BnaFT.A7, BnaFTC6m and BnaFTA7m* driven by *(C+A)-FTp*. T The number of independent T1 plants used for analysis is indicated above the x-axis. (A, B, D) Flowering time was scored as total leaf number. Box plots show the medians as vertical lines; box limits indicate the 25th and 75th percentiles. Different symbols from a to h above the plots indicate significant differences (P < 0.05, two-way analysis of variance (ANOVA) with Tukey’s multiple comparison test).

The florigenic BnaFT.A7 and the non-florigenic BnaFT.C6 of cultivar ZS11 differ at 5 amino-acid residues, of which none correspond to five FT residues (D17, V18, V70, S76, and R83) previously identified as important for intercellular trafficking (Yoo *et al*., 2013; Endo *et al*., 2018) (Supplemental Figure 4). The carboxy-terminus is important for FT function, which is abolished by the deletion of the last nine amino acid residues (Kim *et al*., 2016). The presence of a cysteine at the last position of BnaFT.C6 instead of the threonine of BnaFT.A7 alters biochemical properties and could cause covalent protein-protein cross-linking under oxidizing conditions (Figure 2D). Modified protein versions carrying a mutual exchange of the last residue (BnaFT.A7m and BnaFT.C6m) were expressed in *ft-10* under the control of *C+A-FTp*. T1 plants transformed with *BnaFT.A7m* showed a significant delay in flowering compared to *BnaFT.A7* transformed lines (Figure 2E). In contrast, *BnaFT.C6m* T1 plants did not flower earlier than *BnaFT.C6* transformed plants. Taken together, the data indicate that a cysteine as the last amino-acid contributed to a reduced florigen function of BnaFT.C6, but was not the only determinant. Comparison of BnaFT.C6 from different cultivars revealed that the threonine to cysteine change at the last amino-acid position was only found in a subset of semi-winter cultivars (Supplemental Figure 5).

### Expression of *FT* homologs across tissues

Although *BnaFT.C6*, *BnaNFT.A7* and *BnaCFT.C4* may not encode fully functional mobile florigens, they induce flowering if transcribed close to the apical meristem. We compared steady-state mRNA levels of *BnaFT*-like genes in different tissues and developmental stages in the cultivar ZS11 grown in LD with and without vernalization (Figure 3). Genomic DNA was used as a common reference point to allow copy-number comparisons of mRNA levels between homologs. Without vernalization, no differences in steady-state mRNA levels were detected between leaf 6 and leaf 8. In the case of vernalized plants, leaf 6 and leaf 8 were fully expanded before and after the plants were moved to and out of the vernalization chamber, respectively. *BnaFT.A7* and *BnaFT.C6* were strongly induced in leaf 8 of vernalized compared to non-vernalized plants, while *BnaFT.A2* was vernalization-responsive but less induced. As previously reported, *BnaFT.C2* was barely expressed, which has been attributed to the presence of several TE insertions at the locus (J., Wang *et al*., 2012). After vernalization, *BnaFT.A2* was expressed at much lower levels than *BnaFT.A7*/*BnaFT.C6* in leaf 8, but became the highest expressed *FT*-like gene in paraclade leaves. Steady-state mRNA levels were similar for all three *FT* orthologs in floral buds, flowers and developing siliques. In contrast, expression of *BnaFT.C2, BnaNFT.A7* and *BnaCFT.C4* was low throughout development; therefore, these *FT*-like genes are unlikely to play a major regulatory role for flowering regulation or inflorescence development under our test conditions.

**Figure 3.**
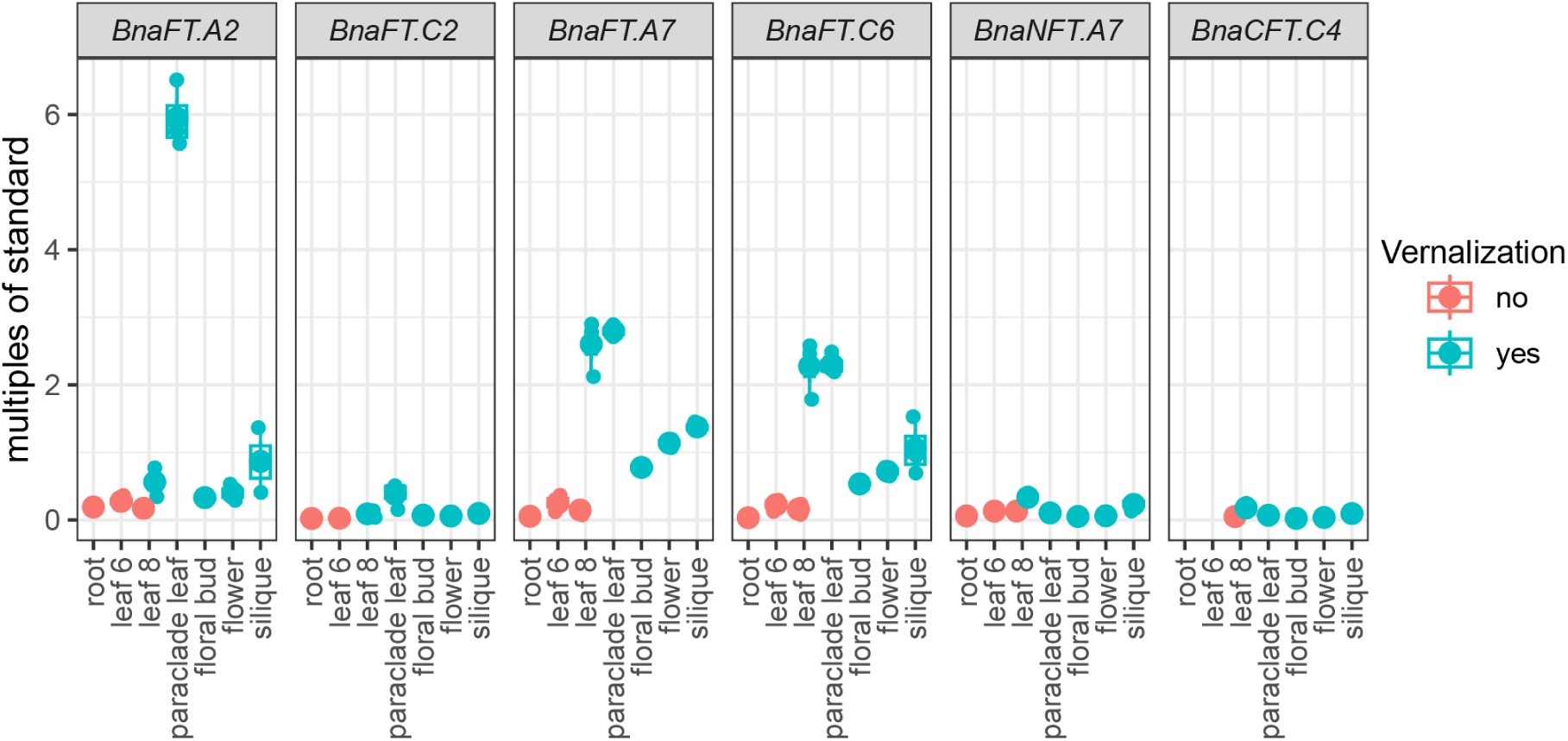
Tissue-specific expression of *B. napus FT* homologs in ZS11. Expression of *B. napus FT* homologs in roots, leaf 6, leaf 8, paraclade leaf, floral bud, flower and siliques. Plants were sown in the greenhouse for 4 weeks under LD conditions, then vernalized for 4 weeks before being transferred back to LD conditions in the greenhouse. Root and leaf 6 and 8 were sampled before vernalization, leaf 8, paraclade leaf, floral bud, flower and siliques were sampled after vernalization at ZT12. Boxes represent inner quartiles of three biological replicates, medians are indicated by vertical lines, single measurements are indicated by dots. *BnaENTH* was used as reference gene, all values were compared to genomic DNA, which was set as 1.

### Conservation of cis-regulatory regions at *B. napus FT* homologs

Previous reports established that the distal *FT* enhancer Block C is conserved at *FT* orthologs between *B. napus* and *A. thaliana* (J., Wang *et al*., 2012; Adrian *et al*., 2010). Here we show that this is also the case for Block E, which is located downstream of the structural gene. *FT* orthologs of *B. oleracea*, *B. rapa* and *S. parvula* showed the presence of all three blocks, while non-orthologous *FT* homologues showed no conservation of the distal blocks and limited conservation at the proximal promoter Block A (Supplementary Figure 6).

In *A. thaliana*, *FT* transcription is dependent on the NF-Y/CO-complex, but also requires canonical NF-Y complexes, that contain NF-YA family proteins instead of CO (Wenkel *et al*., 2006; Siriwardana *et al*., 2016; Kumimoto *et al*., 2010; Kumimoto *et al*., 2008). The effect of NF-Y is likely mediated by cognate CCAAT-boxes located at the distal enhancers Block C and Block E (Zicola *et al*., 2019; Cao *et al*., 2014; Siriwardana *et al*., 2016). A CCACA-motif as high-affinity binding site of the CO-complex is found three times at Block A; a CCACT-motif that still binds the CO-complex with fair affinity is found twice (Gnesutta *et al*., 2017) (Figure 4A). The function of three distinct CCACA-motifs, named P1, P2 and CORE2, has been experimentally validated (Tiwari *et al*., 2010; Adrian *et al*., 2010). In *B. napus*, CO-binding motifs P2 and CORE2 are present at *BnaFT.A2/BnaFT.C2,* P1 is a CCACT-motif in *BnaFT.A7*, *BnaFT.A2*, *BnaFT.C2* and *SpFT*, while an additional C to A change creates a CCAAT-motif at the same position in *BnaFT.C6*. A second change from CCACT-to CCAAT-motif is common to *BnaFT.C6*, *BnaFT.A7* and *SpFT*. *BnaFT.C6* contains a third, private CCAAT-motif close to the transcription start site. A recent structural study of the CO-complex with target DNA indicated that only the CACA core of the first motif was involved in protein-DNA interactions (Lv *et al*., 2021). Two additional CACA-motifs can be identified at Block A of which one is present in all *BnaFT* paralogs and *SpFT*. In sum, CO-binding CCACA-motifs are observed in block A in a decreasing trend from *BnaFT.C6* to *BnaFT.A7*, *SpFT*, *BnaFT.A2/BnaFT.C2* and *FT*, while a trend towards more NF-Y-binding CCAAT-motifs is observed in the opposite order. Notably, the trend to more promoter-located CCAAT-motifs correlated with higher expression levels in leaves.

**Figure 4.**
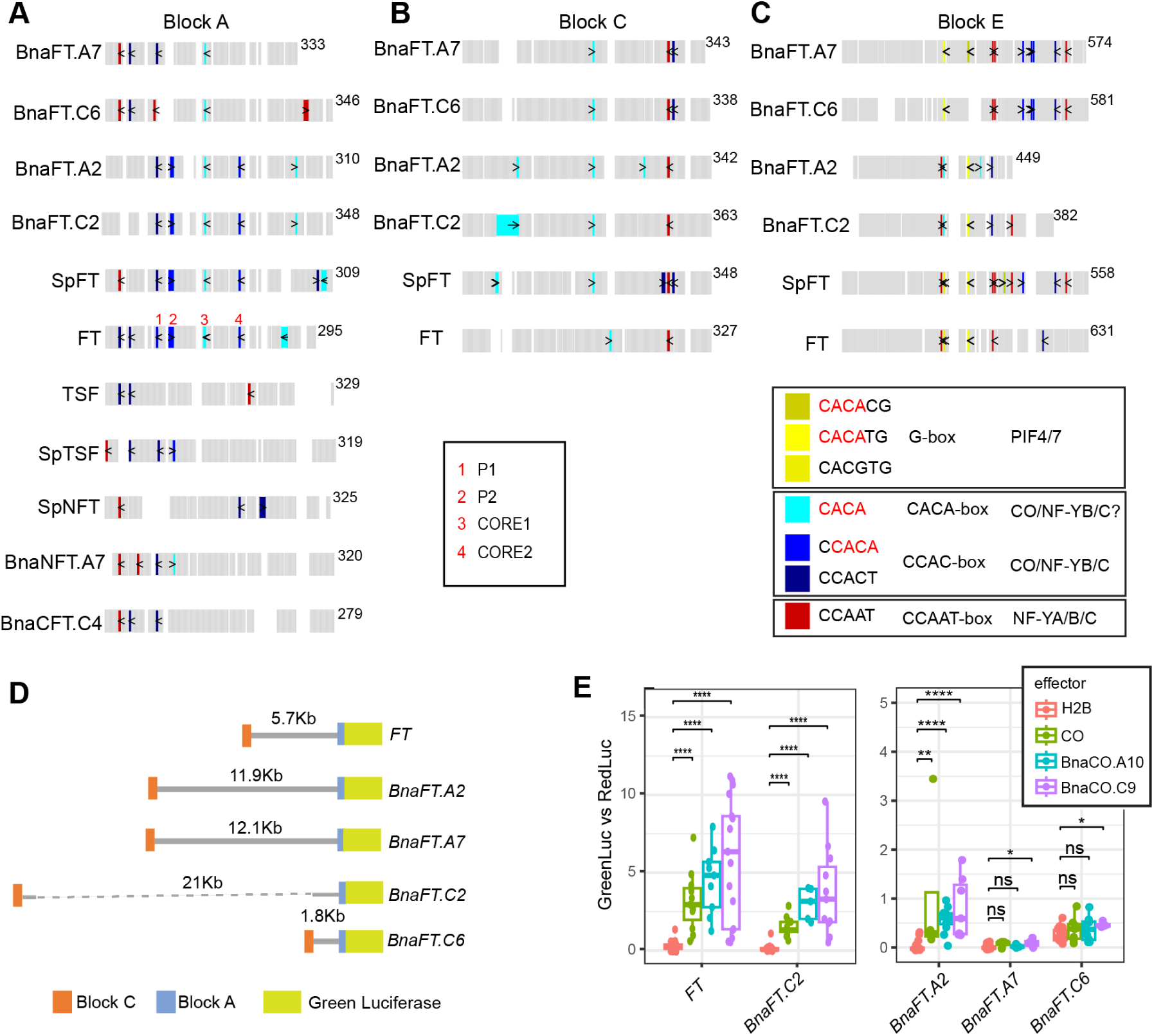
Conservation of the cis-regulatory code at *B. napus FT* homologs. Schematic diagram showing the distribution of motifs within conserved blocks A **(A)**, C **(B)** and E **(C)** for *FT-like* genes from *A. thaliana, S. parvula* and *B. napus*. Gray boxes indicate aligned sequences, white spaces indicate alignment gaps. Cis-elements are marked in different colors as indicated in the common legend. The length of the unaligned sequences is shown after each alignment block. Red numbers above the position of motifs at *FT* block A indicate previously characterized functional motifs. (**D)** Scheme of the promoter region of *B. napus FT* genes fused to LUC68. (**E)** Promoter-reporter transactivation assay in infiltrated tobacco leaves. Agrobacteria transformed with a binary vector containing promoter Firefly green shifted luciferase fusions were co-infiltrated with cultures expressing red shifted luciferase and either CO, BnaCO.A10, BnaCO.C9 or H2B under the control of the CaMV 35S promoter. Luciferase activity was detected with the help of a CCD camera through selective filters. Boxplots show relative green versus red luciferase signal after filter leakage correction from 4 to 6 independent infiltrations of three different leaves, the median is indicated as line, data-points as dots. Statistical analysis was performed using the non-parametric Wilcocxon signed rank test with FDR correction.

At Block C, an important CCAAT-motif is conserved in all *FT* orthologs (Figure 4B), but at the downstream enhancer Block E, *SpFT*, *BnaFT.C6* and *BnaFT.A7* share a CCACA-motif that is not present at the other genes; furthermore, *BnaFTA7* and *BnaFTC6* have evolved two additional CCACA sites as candidates for binding a CO-complex (Figure 4C). In contrast, the first CCAAT motif has been lost at these homeologs.

To test whether the variation in *cis*-motifs at *B. napus FT* promoters had an impact on gene activation by CO, we cloned upstream sequence of *FT* and *B. napus* orthologs to a binary reporter vector with green shifted Luciferase that allowed us to test transactivation by *B. napus* CO homologs in transiently infiltrated tobacco leaves using a co-infiltrated red shifted luciferase for normalization. The distance between regulatory blocks differs greatly in *B.napus* and we could not amplify the 21 Kb promoter region of *BnaFT.C2;* however, we could amplify and fuse the regions corresponding to Block C and Block A (*C+A-BnaFT.C2*) (Figure 4D). We tested ca. 12Kb long *BnaFT.A2* and *BnAFT.A7* promoters, 2Kb long *BnaFT.C6* and *C+A-BnaFT.C2* promoters and a 5.7Kb long *FT* promoter for reporter gene induction in response to *CO*, *BnaCO.A10* and *BnaCO.C9* controlled by the *CaMV 35S* promoter. We found that promoters of *FT*, *BnaFT.A2* and *C+A-BnaFT.C2* were activated in the presence of COs while the homeolog pair *BnaFT.A7/BnaFT.C6* was not or significantly less induced (Figure 4E).

In sum, *BnaFT.A2* and *C+A-BnaFT.C2*, which show conservation of putative CO binding CCACA-motifs at the proximal promoter are responsive to CO homologs in transient assays, while homeologs *BnaFT.A7/BnaFT.C6* respond poorly to transactivation, which corresponds to the absence of conserved CO-binding motifs.

### Diurnal expression pattern of *B. napus FT* homologs in response to photoperiod and vernalization

Since the homeologs Bna*FT.A2/BnaFT.C2* and *BnaFT.A7/BnaFT.C6* showed remarkable differences in the distribution of cis motifs across regulatory blocks, we compared their transcript levels across time-series of plants grown in SD and LD photoperiod both before and after vernalization. The analysis confirmed that *BnaFT.A2*, *BnaFT.A7* and *BnaFT.C6* respond to vernalization; furthermore, expression was induced by LD versus SD conditions, with a strong synergism between LD and vernalization signals (Figure 5). The expression of *BnaFT.C2* was low (approximately 20 and 100 times less than *BnaFT.A2* and *BnaFT.C6,* respectively) but essentially followed the same induction pattern. In the case of non-*FT* orthologs, *BnaCFT.C4* mirrored the vernalization response and the diurnal expression pattern of *BnaFT* genes, at a level 10 times lower than even *BnaFT.C2*. *BnaNFT.A7* was only notably expressed at the beginning of LDs before vernalization. Overall, apart from *BnaNFT.A7*, *FT*-like genes shared an expression minimum at ZT8 and high expression throughout the night until the next morning (Figure 5 A and B).

**Figure 5.**
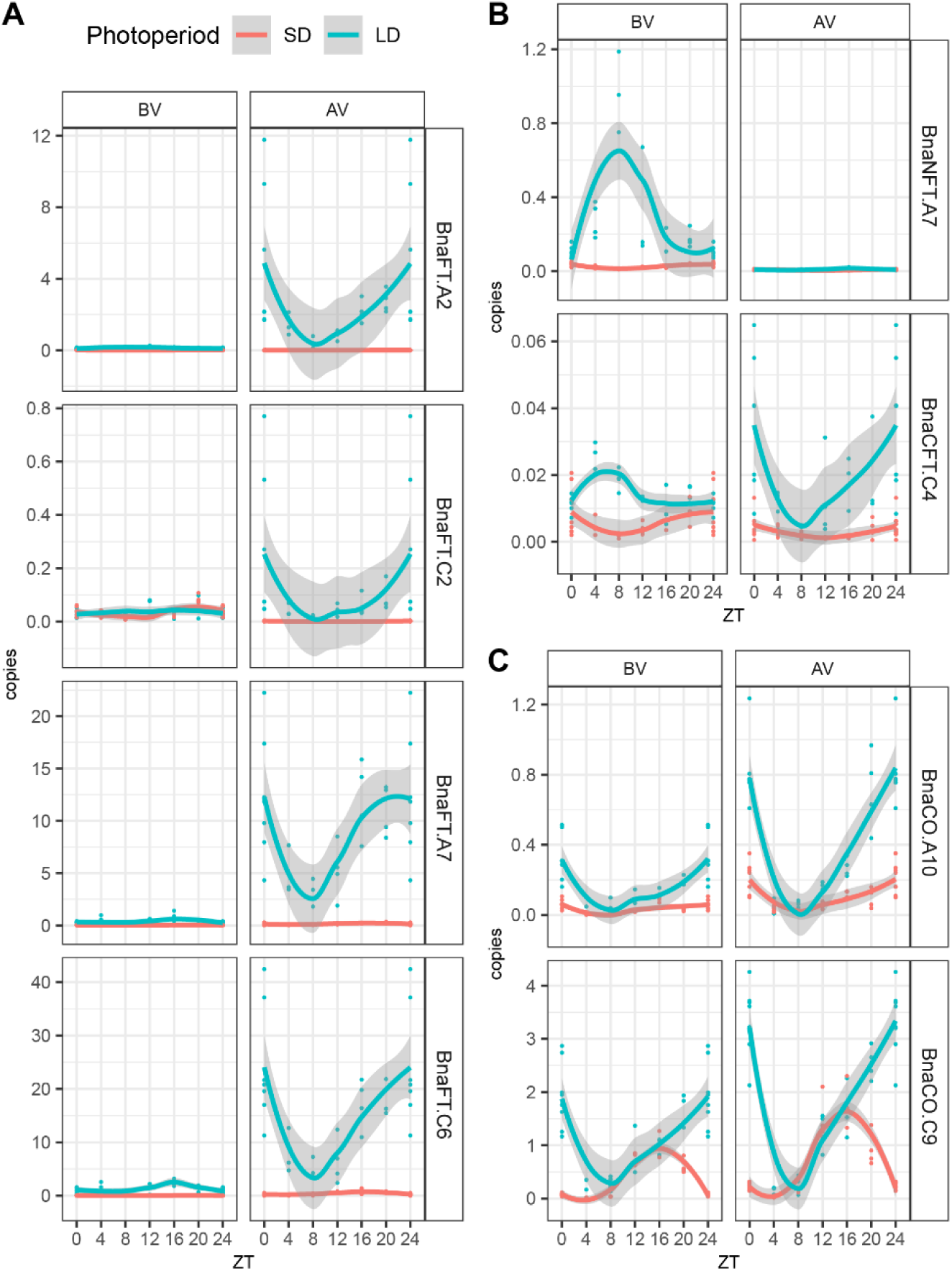
Diurnal expression of *B.* napus *FT* and *CO* homologs in ZS11. **(A)** Diurnal expression of *BnaFT.A2*, *BnaFT.C2*, *BnaFT.A7*, *BnaFT.C6*. **(B)** Diurnal expression of *BnaNFT.A7* and *BnaCFT.C4.* (**C)** Diurnal expression of *BnaCO.A10* and *BnaCO.C9.* Material from leaf 6 and leaf 8 were sampled every 4 h from ZS11 plants grown in SD and LD photoperiod in growth chambers at 22°C before (BV) and after (AV) vernalization, respectively. Time is plotted in reference to Zeitgeber (light on). Expression levels were analysed by gene-specific qRT-PCR using four independent biological replicates using genomic DNA as common reference point. *BnaENTH* was used to correct for differences in cDNA amount. Values for ZT0 and ZT24 were pooled at both time points. Data points are indicated as dots, lines were generated using a loess smoothing function, grey shapes indicate ± standard error.

### Photoperiod-dependent regulation of putative *B. napu*s *FT* activators

Differences in the arrangement of cis-motif for NF-Y/CO-and NF-Y complexes between regulatory regions did not abolish the dependence of *BnaFT* genes on LD photoperiod. In *A. thaliana*, the accumulation of CO protein in the second half of long days largely explains LD dependency of *FT* expression, but it is not yet addressed if this model applies to other Brassicaceae. We measured steady-state transcript levels of *BnaCO.A10* and *BnaCO.C9* in leaves throughout SD and LDs before and after vernalization (Figure 5C). As previously described, *BnaCO.C9* mRNA accumulated at higher levels than *BnaCO.A10* (Q., Jin *et al*., 2021). Both genes showed higher expression in LD vs SD, with similar trends before and after vernalization. As observed for *BnaFT*-like genes, transcript levels of both *BnaCO* genes showed a trough at ZT8 in LD. For *BnaCO.A10,* the general pattern of transcript accumulation was similar in SD and LD, showing an increase during the dark and at dawn. For *BnaCO.C9*, steady-state levels declined 8 h after the onset of the dark period in SD corresponding to the onset of dark in LD, but transcript levels continued to increase in LD during the dark period. Under the assumption that CO proteins in *B. napus* require light for stabilization, only *BnaCO.A10* transcripts accumulating in the morning would contribute to a NF-Y/CO complex in the morning in SD, while both COs could contribute to early morning and late afternoon NF-Y/CO complexes in LD.

To assess whether NF-Y components could also limit *FT* gene expression in *B. napus*, we generated RNAseq libraries of a diurnal time-series of vernalized ZS11 plants grown in LD and SD photoperiod. Of the annotated 95560 nuclear *B. napus* genes, 47198 were considered as expressed (at least 25 transcripts per million (TPM) summed across all samples) (Supplemental Data 2). After applying the Rhythmicity Analysis Incorporating non-Parametric methods (RAIN) pipeline with a FDR <0.05, 33714 genes (71,4%) scored as diurnally expressed, with a slightly higher number in SD vs LD (29596 vs 21757). Most genes were detected as diurnally expressed in both photoperiods (52.3%), but more genes were SD-than LD-specific (35.5% vs 12.2%, respectively) (Figure 6A, Supplemental Data 3).

**Figure 6.**
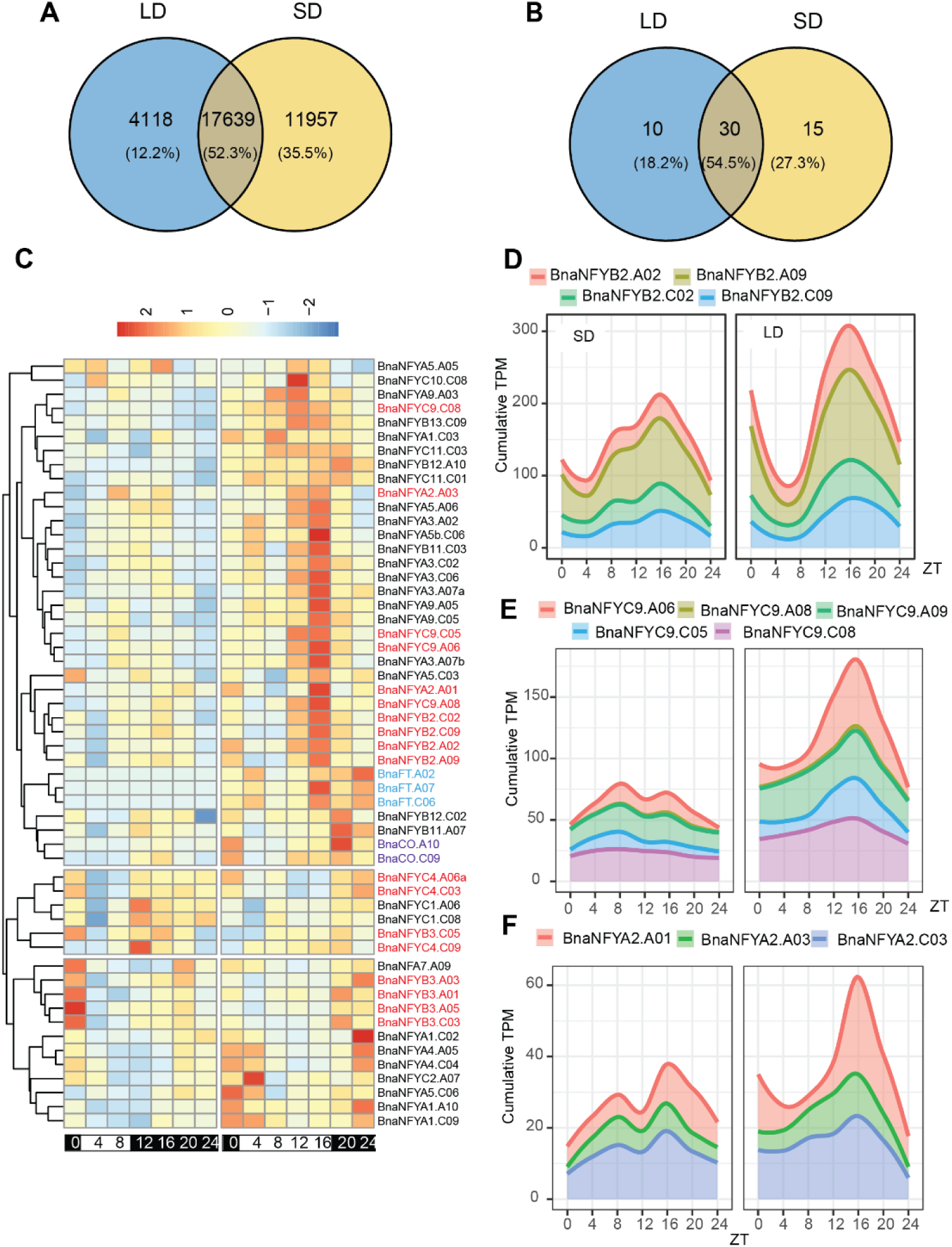
Diurnal expression of *B.* napus NF-Y homologs. **(A)** Venn diagram showing rhythmically expressed genes in LD and SD photoperiod. (**B)** As A) for NF-Y genes. **(C)** Heatmap of rhythmically expressed *B. napus* NF-Y genes after hierarchical clustering based on expression patterns. Expression data in TPM were normalized across all samples per gene, clustering was performed using the Manhattan distance metric. Colours indicate *B. napus* homologs of *NF-Y*s involved in *FT* regulation (red), *BnaFT*s (blue) and *BnaCO*s (purple). (**D)** Cumulative sum plot of TPM values in SD (left panel) and LD (right panel) for all *BnaNFYB2* genes. (**E)** as D) for *BnaNF-YC9* genes. (**F)** as D) for *BnaNF-YA2* genes.

We re-annotated *B. napus* NF-Y genes based on synteny to *A. thaliana* homologs and removed four genes that were annotated as NF-Y genes but that likely encoded by pseudogenes due to a partial protein prediction and/or frame-shift mutations (Supplemental Table 4, Supplemental Figure 6). We found that 78 of 106 NF-Y genes were expressed, of which 55 showed diurnal changes in their transcript levels. The fraction of diurnally expressed NF-Y genes is as expected and although a slightly greater than expected number of NF-Y genes showed LD-specific rhythmicity, this was not statistically significant (Figure 6B). Hierarchical clustering of diurnally expressed *B. napus* NF-Y genes based on their transcript accumulation patterns (normalized across all samples) identified 3 major clades (Figure 6C). A clade containing 29 NF-Y genes was expressed at higher level in LD versus SD and showed a marked peak of expression in the critical afternoon light period from ZT12 to ZT16 in LD. The clade also included *BnaFT* and CO genes that had been added to the clustering analysis; however, expression of these genes extended to the dark phase and/or showed later peak values than ZT16.

In *A. thaliana*, NF-Y paralogs with a critical role in *FT* expression were identified by their phloem-specific expression and subsequently tested by reverse genetic approaches (Kumimoto *et al*., 2010; Siriwardana *et al*., 2016; Kumimoto *et al*., 2008). Although the tissue-specific expression pattern of NF-Y genes in *B. napus* is unknown, it is noteworthy that 4 of 4 NF-YB2 homologs, 4 of 5 NF-YC9 homologs and 2 of 3 NF-YA2 homologs, all orthologs of photoperiod-responsive NF-Ys, are part of the expression cluster with an LD-specific peak at ZT16. *B. napus* homeologs are likely to act redundantly and we therefore prepared cumulative plots that consider the expression of all *B. napus* NF-YB2, NF-YC9 and NF-YA2 genes (Figure 6 D-F). If the sum of the expression of all homeologs is considered, a marked peak of expression at ZT16 in LD and an overall higher expression in LD vs SD was detected. In sum, NF-Y genes that are candidates for photoperiod-dependent induction of *FT*-like genes could strongly promote *FT* expression at the end of LD.

## Discussion

Land plants share a recurrent history of whole genome duplications (WGD) through polyploidisation and subsequent retraction of the coding genome (Panchy *et al*., 2016). In some cases, homeologs may be retained if increased gene dosage of redundant genes confers a fitness advantage, but more often retention is linked to partial redundancy either through the evolution of divergent expression patterns or through preferred interactions with other protein partners.

*B. napus* is a particularly rewarding case for study, combining two paleontological events with different outcomes with a recent hybridisation event that has been partially shaped by human selection for crop traits (Lu *et al*., 2019; Parkin *et al*., 2005).

Analysis of the genome of the genus Brassica suggests that one of the paleo-duplications resulted in unequal retention of homeologs, while for the second event, two fairly co-linear genomes can still be recognized (Parkin *et al*., 2005). Only two *FT* syntenic genes are conserved per Brassica genome lineage, indicating a loss of the copy of the weaker genome from the first polyploidisation event. *NFT* and *CFT* are not survivors of the weak genome as syntenic copies can be detected in *S. paruvula,* a sister genus of *Brassica* that did not undergo whole genome duplication (Figure 1) (Cheng *et al*., 2012). Differences in expression may have contributed to the maintenance of both *FT* paralogs. In the case of the Brassica A genome, both paralogs are expressed; however, expression seems specialised as *BnaFT.A7* and *BnaFT.A2* show higher expression in rosette and paraclade leaves, respectively (Figure 3). In the case of the C-genome, *BnaFT.C2* is barely expressed, probably due to the presence of TE insertions that are common to *B. napus* but not to all *B. oleracea* varieties (J., Wang *et al*., 2012). Since *BnaFT.C2* encodes a fully functional florigen (Figure 2), the presence of a second *FT*, even if only marginally expressed, may be advantageous under certain conditions and therefore retained. Similarly, circumstances causing the induction of *BnaNFT.C6* and *BnaCFT.C4* could influence flowering and convey an advantage; however, *BnaNFT.C6* homeologs have not survived as functional genes (Figure 1). Similarly, *BnaFT.C6*, the second *FT* of the C-genome may be impaired or at least attenuated for its florigen function (Figure 2). The amino acid change from threonine to cysteine, which contributed to the reduced systemic function, is not found in all *B. napus* accessions (Supplemental Figure 4). Further studies are required to clarify whether the mutation can be linked to a flowering QTL. In the case of the semi-winter *B. napus* variant ZS11, both *FT* genes originating from the C-genome seem compromised in their function.

*A. thaliana FT* regulation depends on an intricate interplay of two distal enhancers with an accessible chromatin structure and a proximal promoter that is inaccessible (Zicola *et al*., 2019). Binding of NF-Y complexes to distal enhancers is thought to somehow increase the probability of NF-Y/CO to bind to the proximal promoter and increase expression; how NF-Y/CO binding is mechanistically linked the induction of transcription is currently unknown. Although the cis-regulatory Blocks were found as conserved at all *B. napus FT* genes, the shift from NF-Y/CO to NF-Y binding sites at the proximal promoter was unexpected and raises a couple of questions (Figure 4). First, can binding of NF-Y simply replace NF-Y/CO causing similar effects leading to the induction of transcripts? Second, if NF-Y binding is sufficient to induce expression, what limits induction of BnaFTA7/BnaFTC6 to LD? Here, we propose that transcriptional regulation of NF-Y genes may contribute to photoperiod control (Figure 6). Alternatively, *B.napus* CO proteins as part of a NF-Y/CO complex binding to the distal enhancer could contribute as limiting factors, but how it could do so is not obvious. Gnesutta et al. determined the binding preferences of the NF-Y/CO complex to DNA (Gnesutta *et al*., 2017). The preferred binding motif for NF-Y/CO (CCACA) is two nucleotide edits away from the CCAAT motif. We found that replacement of motifs happened at conserved in the proximal promoter and propose that an intermediate CCACT motif that can still be bound (or already be bound) by NF-Y/CO served as stepping stone during the motif change (Figure 4).

In sum, *B. napus* photoperiod dependent regulation of *FT*-like genes is a case study for the consequence of gene duplications, indicating gene retraction through loss-of-function, homeolog retention partially explained by differences transcriptional regulation and possibly variation between cultivated varieties due to changes in protein function.

## Data availability

RNAseq datasets generated and analysed in the current study are available at the European Nucleotide archive under the accession number project accession number PRJEB82128.

## Acknowledgements

JW and ND received funding from the Chinese Scholarship Council. FT, HRZ and PT are supported by a Core Grant from the Max Planck Society. FT also receives funding from the DFG through the Cluster of Excellence CEPLAS (EXC 2048/1 Project ID: 390686111).

## Author contributions

JJW and FT outlined, created figures and wrote the manuscript outline. JJW and PT performed all experimental work, ND provided the FDp-FT-pER8 construct and helped with cloning, FT and HRZ contributed with guidance and critical discussion. JJW and FT performed all data analysis. JJW, FT, HRZ and JW were involved in revising the manuscript.

## References

Abe, M., Kobayashi, Y., Yamamoto, S., et al. (2005) FD, a bZIP protein mediating signals from the floral pathway integrator FT at the shoot apex. Science, 309, 1052–1056.

Adrian, J., Farrona, S., Reimer, J.J., Albani, M.C., Coupland, G. and Turck, F. (2010) cis-Regulatory elements and chromatin state coordinately control temporal and spatial expression of FLOWERING LOCUS T in Arabidopsis. Plant Cell, 22, 1425–1440.

Andrés, F. and Coupland, G. (2012) The genetic basis of flowering responses to seasonal cues. Nat. Rev. Genet., 13, 627–639.

Bratzel, F. and Turck, F. (2015) Molecular memories in the regulation of seasonal flowering: from competence to cessation. Genome Biol., 16, 192.

Cao, S., Kumimoto, R.W., Gnesutta, N., Calogero, A.M., Mantovani, R. and Holt, B.F. (2014) A distal CCAAT/NUCLEAR FACTOR Y complex promotes chromatin looping at the FLOWERING LOCUS T promoter and regulates the timing of flowering in Arabidopsis. Plant Cell, 26, 1009–1017.

Chalhoub, B. (2014) Early allopolyploid evolution in the post-Neolithic Brassica napus oilseed genome (vol 348, 1260782, 2014). *Science*, 345, 1255–1255.

Cheng, F., Mandáková, T., Wu, J., Xie, Q., Lysak, M.A. and Wang, X. (2013) Deciphering the diploid ancestral genome of the Mesohexaploid Brassica rapa. Plant Cell, 25, 1541–1554.

Cheng, F., Wu, J., Fang, L. and Wang, X. (2012) Syntenic gene analysis between Brassica rapa and other Brassicaceae species. Front. Plant Sci., 3, 198.

Clough, S.J. and Bent, A.F. (1998) Floral dip: a simplified method for *Agrobacterium*-mediated transformation of *Arabidopsis thaliana*. Plant J., 16, 735–743.

Dobin, A., Davis, C.A., Schlesinger, F., Drenkow, J., Zaleski, C., Jha, S., Batut, P., Chaisson, M. and Gingeras, T.R. (2013) STAR: ultrafast universal RNA-seq aligner. Bioinformatics, 29, 15–21.

Endo, M., Yoshida, M., Sasaki, Y., et al. (2018) Re-Evaluation of Florigen Transport Kinetics with Separation of Functions by Mutations That Uncouple Flowering Initiation and Long-Distance Transport. Plant Cell Physiol., 59, 1621–1629.

Fernández, V., Takahashi, Y., Le Gourrierec, J. and Coupland, G. (2016) Photoperiodic and thermosensory pathways interact through CONSTANS to promote flowering at high temperature under short days. Plant J., 86, 426–440.

Fornara, F., Panigrahi, K.C.S., Gissot, L., Sauerbrunn, N., Rühl, M., Jarillo, J.A. and Coupland, G. (2009) Arabidopsis DOF transcription factors act redundantly to reduce CONSTANS expression and are essential for a photoperiodic flowering response. Dev. Cell, 17, 75–86.

Galvāo, V.C., Fiorucci, A.-S., Trevisan, M., Franco-Zorilla, J.M., Goyal, A., Schmid-Siegert, E., Solano, R. and Fankhauser, C. (2019) PIF transcription factors link a neighbor threat cue to accelerated reproduction in Arabidopsis. Nat. Commun., 10, 4005.

Gnesutta, N., Kumimoto, R.W., Swain, S., Chiara, M., Siriwardana, C., Horner, D.S., Holt, B.F. and Mantovani, R. (2017) CONSTANS Imparts DNA Sequence Specificity to the Histone Fold NF-YB/NF-YC Dimer. Plant Cell, 29, 1516–1532.

Guo, Y., Hans, H., Christian, J. and Molina, C. (2014) Mutations in single FT-and TFL1-paralogs of rapeseed (Brassica napus L.) and their impact on flowering time and yield components. Front. Plant Sci., 5, 282.

Hatje, K., Keller, O., Hammesfahr, B., Pillmann, H., Waack, S. and Kollmar, M. (2011) Cross-species protein sequence and gene structure prediction with fine-tuned Webscipio 2.0 and Scipio. BMC Res. Notes, 4, 265.

Imaizumi, T., Schultz, T.F., Harmon, F.G., Ho, L.A. and Kay, S.A. (2005) FKF1 F-box protein mediates cyclic degradation of a repressor of CONSTANS in Arabidopsis. Science, 309, 293–297.

Jaeger, K.E. and Wigge, P.A. (2007) FT protein acts as a long-range signal in Arabidopsis. Curr. Biol., 17, 1050–1054.

Jang, S., Marchal, V., Panigrahi, K.C.S., Wenkel, S., Soppe, W., Deng, X.-W., Valverde, F. and Coupland, G. (2008) Arabidopsis COP1 shapes the temporal pattern of CO accumulation conferring a photoperiodic flowering response. EMBO J., 27, 1277–1288.

Jin, Q., Gao, G., Guo, C., et al. (2022) Transposon insertions within alleles of BnaFT.A2 are associated with seasonal crop type in rapeseed. Theor. Appl. Genet., 135, 3469–3483.

Jin, Q., Yin, S., Li, G., et al. (2021) Functional homoeologous alleles of CONSTANS contribute to seasonal crop type in rapeseed. Theor. Appl. Genet., 134, 3287–3303.

Jin, S., Nasim, Z., Susila, H. and Ahn, J.H. (2021) Evolution and functional diversification of FLOWERING LOCUS T/TERMINAL FLOWER 1 family genes in plants. Semin. Cell Dev. Biol., 109, 20–30.

Kagale, S., Robinson, S.J., Nixon, J., et al. (2014) Polyploid evolution of the Brassicaceae during the Cenozoic era. Plant Cell, 26, 2777–2791.

Kiełbasa, S.M., Wan, R., Sato, K., Horton, P. and Frith, M.C. (2011) Adaptive seeds tame genomic sequence comparison. Genome Res., 21, 487–493.

Kim, S.-J., Hong, S.M., Yoo, S.J., Moon, S., Jung, H.S. and Ahn, J.H. (2016) Post-Translational Regulation of FLOWERING LOCUS T Protein in Arabidopsis. Mol. Plant, 9, 308–311.

Kumar, S.V., Lucyshyn, D., Jaeger, K.E., Alós, E., Alvey, E., Harberd, N.P. and Wigge, P.A. (2012) Transcription factor PIF4 controls the thermosensory activation of flowering. Nature, 484, 242–245.

Kumimoto, R.W., Adam, L., Hymus, G.J., Repetti, P.P., Reuber, T.L., Marion, C.M., Hempel, F.D. and Ratcliffe, O.J. (2008) The Nuclear Factor Y subunits NF-YB2 and NF-YB3 play additive roles in the promotion of flowering by inductive long-day photoperiods in Arabidopsis. Planta, 228, 709–723.

Kumimoto, R.W., Zhang, Y., Siefers, N. and Holt, B.F. (2010) NF-YC3, NF-YC4 and NF-YC9 are required for CONSTANS-mediated, photoperiod-dependent flowering in Arabidopsis thaliana. Plant J., 63, 379–391.

Laubinger, S., Marchal, V., Le Gourrierec, J., et al. (2006) Arabidopsis SPA proteins regulate photoperiodic flowering and interact with the floral inducer CONSTANS to regulate its stability. Development, 133, 3213–3222.

Liu, L., Adrian, J., Pankin, A., Hu, J., Dong, X., Korff, M. von and Turck, F. (2014) Induced and natural variation of promoter length modulates the photoperiodic response of FLOWERING LOCUS T. Nat. Commun., 5, 4558.

Liu, Y., Li, X., Li, K., Liu, H. and Lin, C. (2013) Multiple bHLH proteins form heterodimers to mediate CRY2-dependent regulation of flowering-time in Arabidopsis. PLoS Genet., 9, e1003861.

Liu, Y., Li, X., Ma, D., Chen, Z., Wang, J.-W. and Liu, H. (2018) CIB1 and CO interact to mediate CRY2-dependent regulation of flowering. EMBO Rep., 19.

Li, B. and Dewey, C.N. (2011) RSEM: accurate transcript quantification from RNA-Seq data with or without a reference genome. BMC Bioinformatics, 12, 323.

Lu, K., Wei, L., Li, X., et al. (2019) Whole-genome resequencing reveals Brassica napus origin and genetic loci involved in its improvement. Nat. Commun., 10, 1154.

Lv, X., Zeng, X., Hu, H., et al. (2021) Structural insights into the multivalent binding of the Arabidopsis FLOWERING LOCUS T promoter by the CO-NF-Y master transcription factor complex. Plant Cell, 33, 1182–1195.

Panchy, N., Lehti-Shiu, M. and Shiu, S.-H. (2016) Evolution of gene duplication in plants. Plant Physiol., 171, 2294–2316.

Parkin, I.A.P., Gulden, S.M., Sharpe, A.G., Lukens, L., Trick, M., Osborn, T.C. and Lydiate, D.J. (2005) Segmental structure of the Brassica napus genome based on comparative analysis with Arabidopsis thaliana. Genetics, 171, 765–781.

Pin, P.A., Benlloch, R., Bonnet, D., Wremerth-Weich, E., Kraft, T., Gielen, J.J.L. and Nilsson, O. (2010) An antagonistic pair of FT homologs mediates the control of flowering time in sugar beet. Science, 330, 1397–1400.

Posé, D., Verhage, L., Ott, F., Yant, L., Mathieu, J., Angenent, G.C., Immink, R.G.H. and Schmid, M. (2013) Temperature-dependent regulation of flowering by antagonistic FLM variants. Nature, 503, 414–417.

Sawa, M., Nusinow, D.A., Kay, S.A. and Imaizumi, T. (2007) FKF1 and GIGANTEA complex formation is required for day-length measurement in Arabidopsis. Science, 318, 261–265.

Schiessl, S., Samans, B., Hüttel, B., Reinhard, R. and Snowdon, R.J. (2014) Capturing sequence variation among flowering-time regulatory gene homologs in the allopolyploid crop species Brassica napus. Front. Plant Sci., 5, 404.

Schiessl, S. (2020) Regulation and Subfunctionalization of Flowering Time Genes in the Allotetraploid Oil Crop Brassica napus. Front. Plant Sci., 11, 605155.

Siriwardana, C.L., Gnesutta, N., Kumimoto, R.W., Jones, D.S., Myers, Z.A., Mantovani, R. and Holt, B.F. (2016) NUCLEAR FACTOR Y, Subunit A (NF-YA) Proteins Positively Regulate Flowering and Act Through FLOWERING LOCUS T. PLoS Genet., 12, e1006496.

Song, J.-M., Guan, Z., Hu, J., et al. (2020) Eight high-quality genomes reveal pan-genome architecture and ecotype differentiation of Brassica napus. Nat. Plants, 6, 34–45.

Sparkes, I.A., Runions, J., Kearns, A. and Hawes, C. (2006) Rapid, transient expression of fluorescent fusion proteins in tobacco plants and generation of stably transformed plants. Nat. Protoc., 1, 2019–2025.

Takagi, H., Hempton, A.K. and Imaizumi, T. (2023) Photoperiodic flowering in Arabidopsis: Multilayered regulatory mechanisms of CONSTANS and the florigen FLOWERING LOCUS T. Plant Commun., 4, 100552.

Tamura, K., Stecher, G. and Kumar, S. (2021) MEGA11: Molecular Evolutionary Genetics Analysis version 11. Mol. Biol. Evol., 38, 3022–3027.

Tang, H., Krishnakumar, V., Zeng, X., et al. (2024) JCVI: A versatile toolkit for comparative genomics analysis. iMeta, 3, e211.

Tang, H., Wang, X., Bowers, J.E., Ming, R., Alam, M. and Paterson, A.H. (2008) Unraveling ancient hexaploidy through multiply-aligned angiosperm gene maps. Genome Res., 18, 1944–1954.

Teotia, S. and Tang, G. (2015) To bloom or not to bloom: role of microRNAs in plant flowering. Mol. Plant, 8, 359–377.

Thaben, P.F. and Westermark, P.O. (2014) Detecting rhythms in time series with RAIN. J. Biol. Rhythms, 29, 391–400.

Tiwari, S.B., Shen, Y., Chang, H.-C., et al. (2010) The flowering time regulator CONSTANS is recruited to the FLOWERING LOCUS T promoter via a unique cis-element. New Phytol., 187, 57–66.

Valverde, F., Mouradov, A., Soppe, W., Ravenscroft, D., Samach, A. and Coupland, G. (2004) Photoreceptor regulation of CONSTANS protein in photoperiodic flowering. Science, 303, 1003–1006.

Wang, J., Hopkins, C.J., Hou, J., Zou, X., Wang, C., Long, Y., Kurup, S., King, G.J. and Meng, J. (2012) Promoter variation and transcript divergence in Brassicaceae lineages of FLOWERING LOCUS T. PLoS ONE, 7, e47127.

Wang, J., Long, Y., Wu, B., Liu, J., Jiang, C., Shi, L., Zhao, J., King, G.J. and Meng, J. (2009) The evolution of Brassica napus FLOWERING LOCUS T paralogues in the context of inverted chromosomal duplication blocks. BMC Evol. Biol., 9, 271.

Wang, Y., Tang, H., Debarry, J.D., et al. (2012) MCScanX: a toolkit for detection and evolutionary analysis of gene synteny and collinearity. Nucleic Acids Res., 40, e49.

Wenkel, S., Turck, F., Singer, K., Gissot, L., Le Gourrierec, J., Samach, A. and Coupland, G. (2006) CONSTANS and the CCAAT box binding complex share a functionally important domain and interact to regulate flowering of Arabidopsis. Plant Cell, 18, 2971–2984.

Wigge, P.A., Kim, M.C., Jaeger, K.E., Busch, W., Schmid, M., Lohmann, J.U. and Weigel, D. (2005) Integration of spatial and temporal information during floral induction in Arabidopsis. Science, 309, 1056–1059.

Wu, D., Liang, Z., Yan, T., et al. (2019) Whole-Genome Resequencing of a Worldwide Collection of Rapeseed Accessions Reveals the Genetic Basis of Ecotype Divergence. Mol. Plant, 12, 30–43.

Yamaguchi, A., Kobayashi, Y., Goto, K., Abe, M. and Araki, T. (2005) TWIN SISTER OF FT (TSF) acts as a floral pathway integrator redundantly with FT. Plant Cell Physiol., 46, 1175–1189.

Yanovsky, M.J. and Kay, S.A. (2002) Molecular basis of seasonal time measurement in Arabidopsis. Nature, 419, 308–312.

Yoo, S.-C., Chen, C., Rojas, M., Daimon, Y., Ham, B.-K., Araki, T. and Lucas, W.J. (2013) Phloem long-distance delivery of FLOWERING LOCUS T (FT) to the apex. Plant J., 75, 456–468.

Zicola, J., Liu, L., Tänzler, P. and Turck, F. (2019) Targeted DNA methylation represses two enhancers of FLOWERING LOCUS T in Arabidopsis thaliana. Nat. Plants, 5, 300– 307.

Zou, J., Mao, L., Qiu, J., et al. (2019) Genome-wide selection footprints and deleterious variations in young Asian allotetraploid rapeseed. Plant Biotechnol. J., 17, 1998–2010.

