## Supplemental_Figures_and_Tables for "Characterization of *FLOWERING LOCUS T* related genes and their putative gene regulatory network in semi-winter *Brassica napus* cultivar Zhongshaung11"

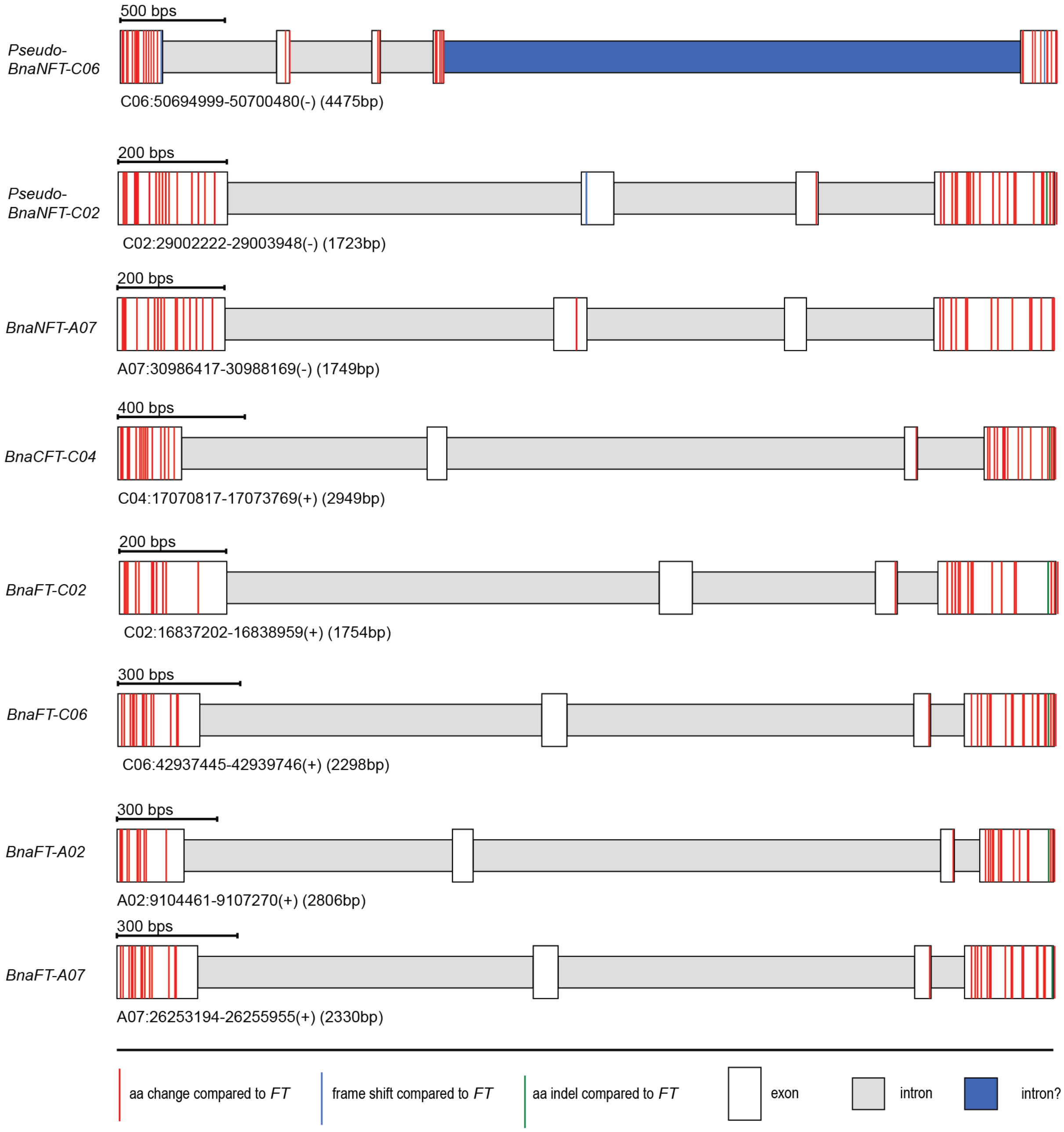


**Supplemental Figure 1. Gene structure of *FT* homologous genes in *B. napus*.** Boxes indicate protein coding regions, red lines indicate aa changes, blue lines frame shifts and green lines in-frame amino-acid indels compared to FT.

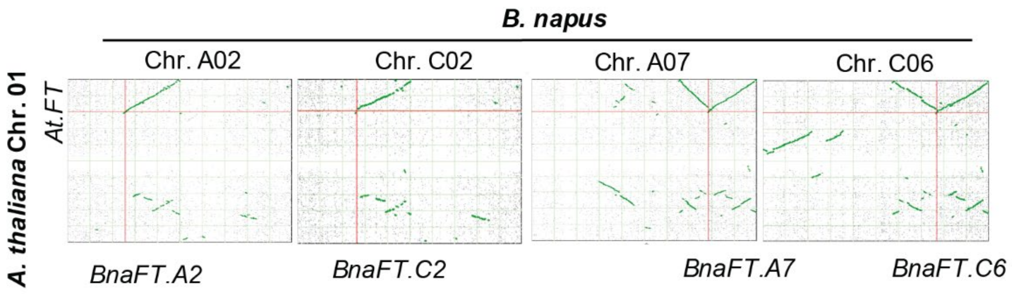


**Supplemental Figure 2. Synteny analysis of *FT* in *A. thaliana* against *B. napus* ZS11 genomic background.**

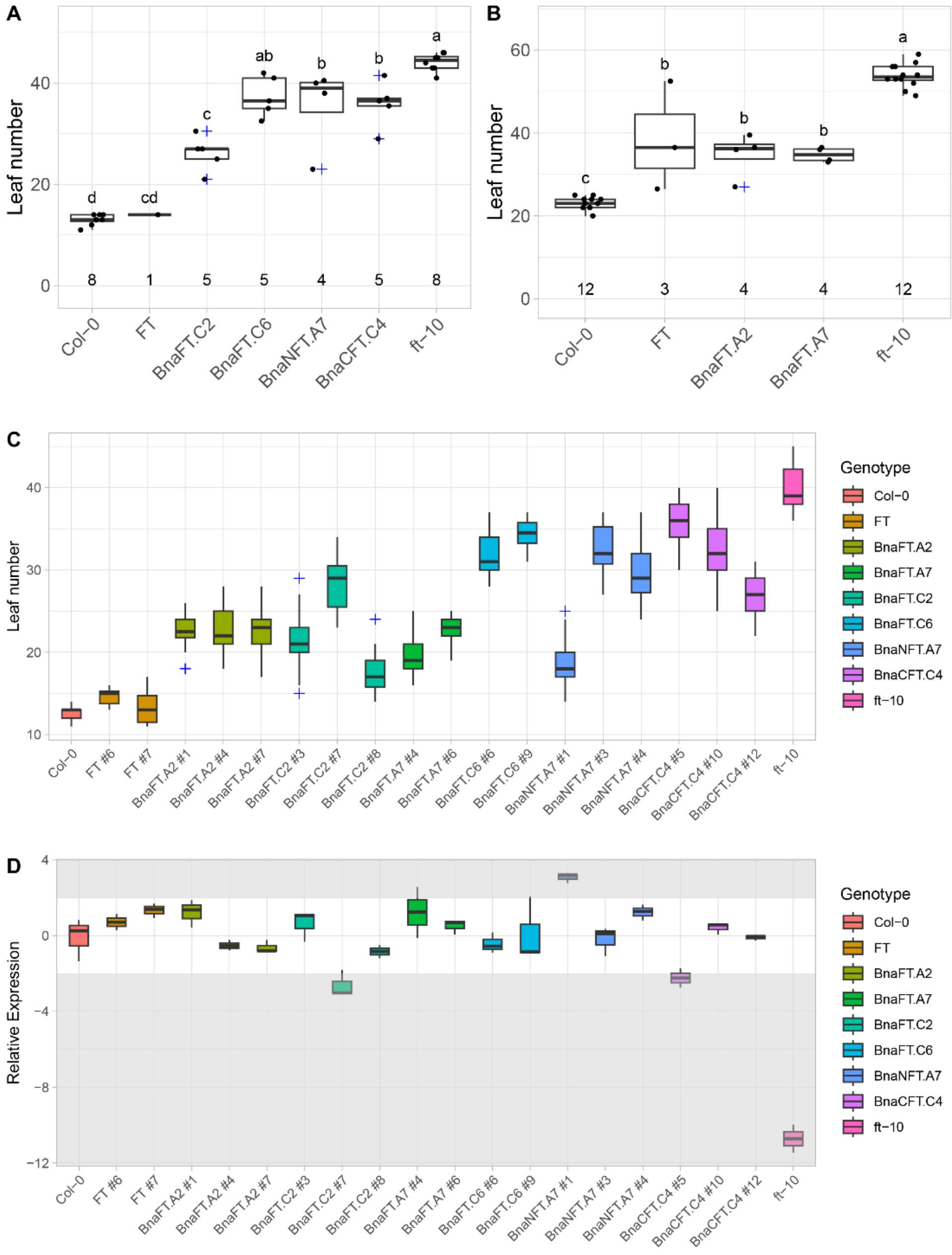


**Supplemental Figure 3. (A) and (B)** Flowering time analysis of independent T2 lines. Flowering time is measured as total leaf number from segregating independent T2 lines grown in LD at 22°C in greenhouse conditions. Values of at least 10 plants from segregating lines were collapsed to their median. Statistical testing was performed by one-factor ANOVA followed by HSD, letters indicate confidence intervals. The number of independent lines is indicated above the category axis within each graph. **(C)** Flowering time analysis of T3 lines from T2 lines showing 3:1 segregation in A and B. **(D)** Analysis of transgene expression in individual T3 lines analysed in (C). Samples for RNA extraction were collected at ZT16 from three biological replicates of 14-day old seedlings grown in LD at 22°C in greenhouse conditions. Expression data were titrated against the plasmids used for transformation using a primer pair against the BAR marker gene to correct for dilution differences. PP2A was used as housekeeping gene, all data were related to the mean of FT expression in Col-0. The white area indicates a band within +/-log2-fold change of 2. Within this area, flowering time appears mostly uncorrelated to expression level, while outliers in the grey shaded area show a corresponding flowering response.

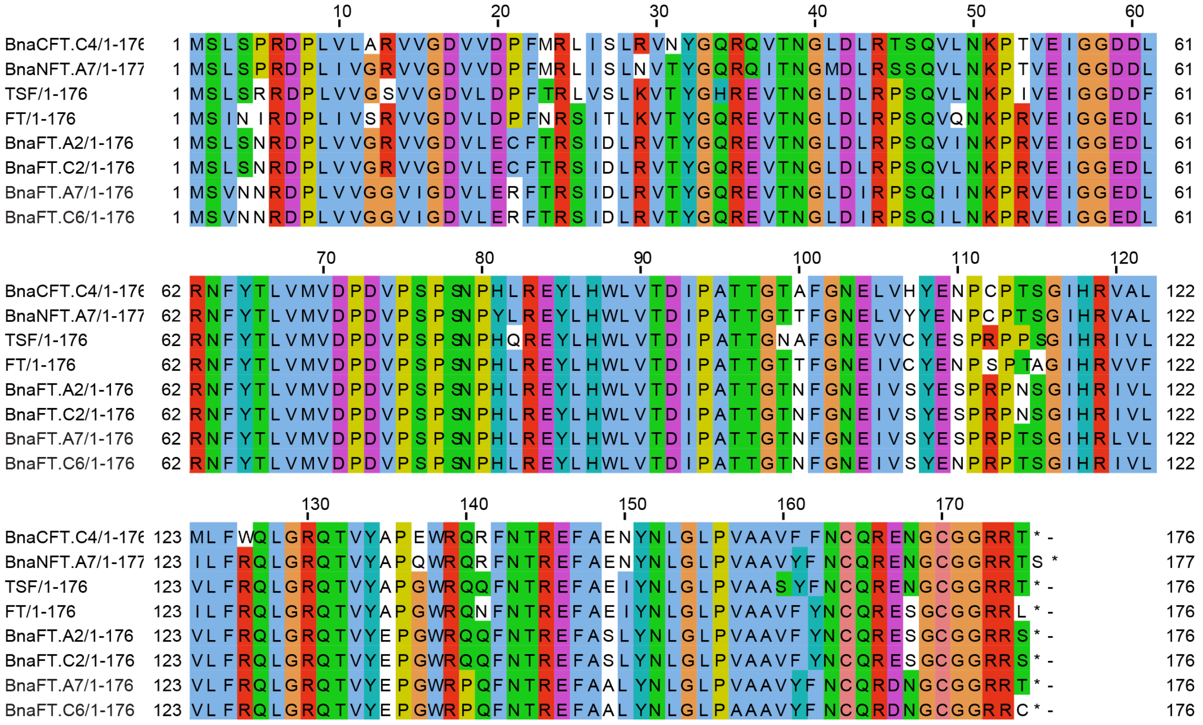

**Supplemental Figure 4. Alignment of FT-like genes from *B. napus* and *A. thaliana*.**

Similar colours indicate conserved amino-acids with similar biochemical properties.

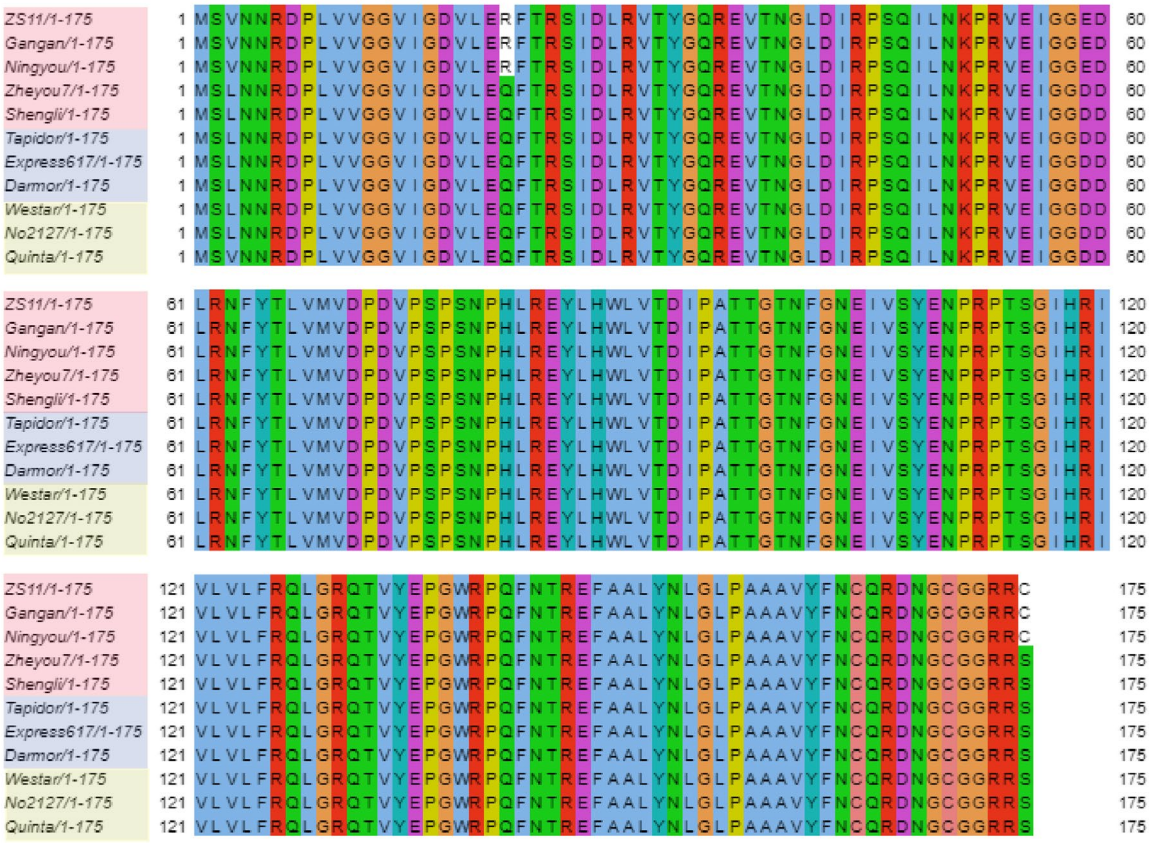

**Supplemental Figure 5. Alignment of BnaFT.C6 from different accessions.** Coloured boxes in indicate semi-winter (red), winter (blue) and spring (yellow) cultivars.

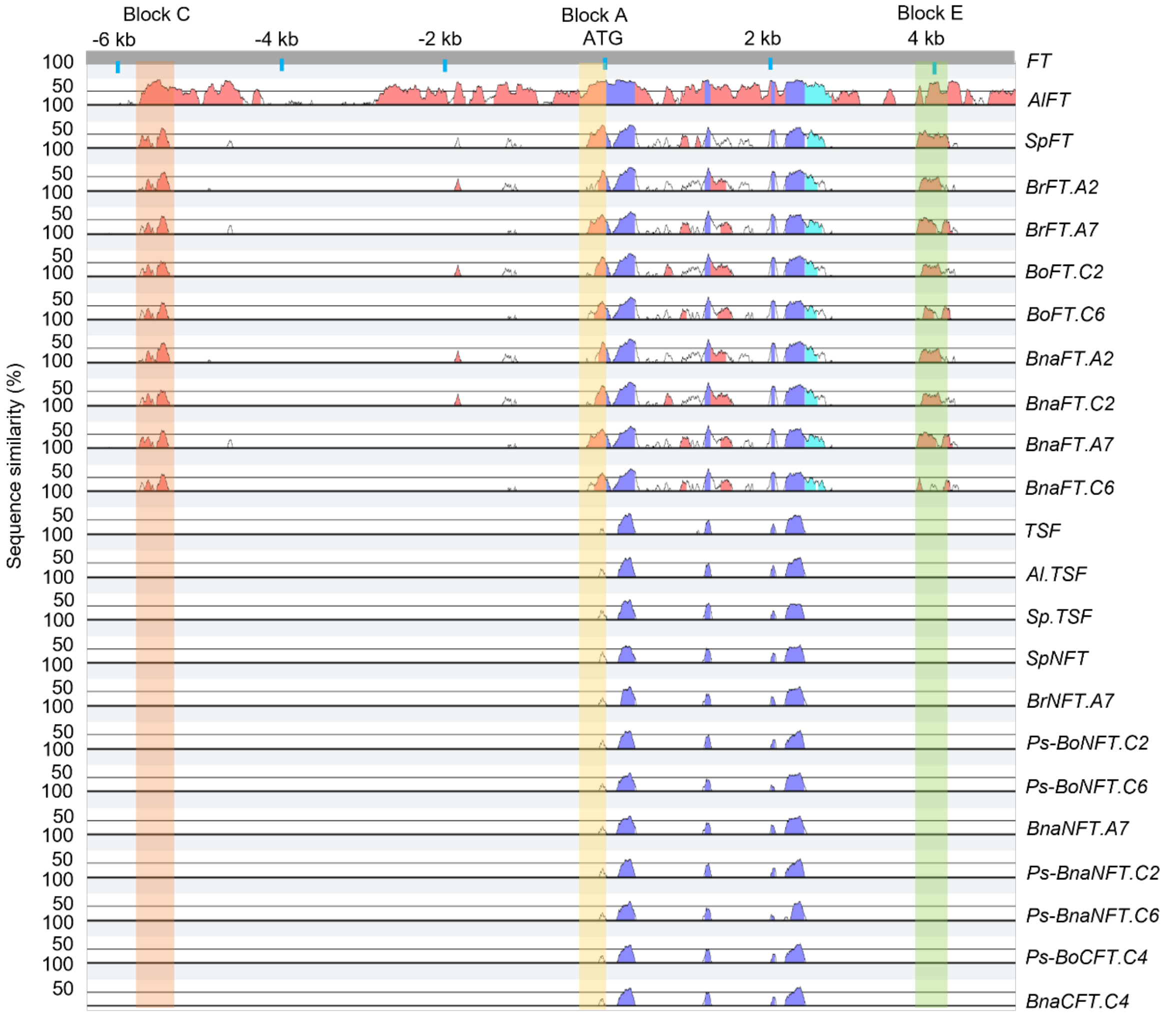


**Supplemental Figure 6. Genome structure alignment analysis.**

Pairwise alignment of genomic sequences of *FT*, *TSF*, *NFT*, *CFT* homologs from different species to the *FT* genomic sequence using mVISTA. The graphical output shows base-pair identity in sliding 100-bp windows in a range of 50% to 100%. The pink regions are "Conserved Non-Coding Sequences" ("CNS"), the dark blue regions are exons, and the light-blue regions are UTRs. Orange, yellow and green boxes indicate the location of conserved Block C, Block A, and Block E, respectively.

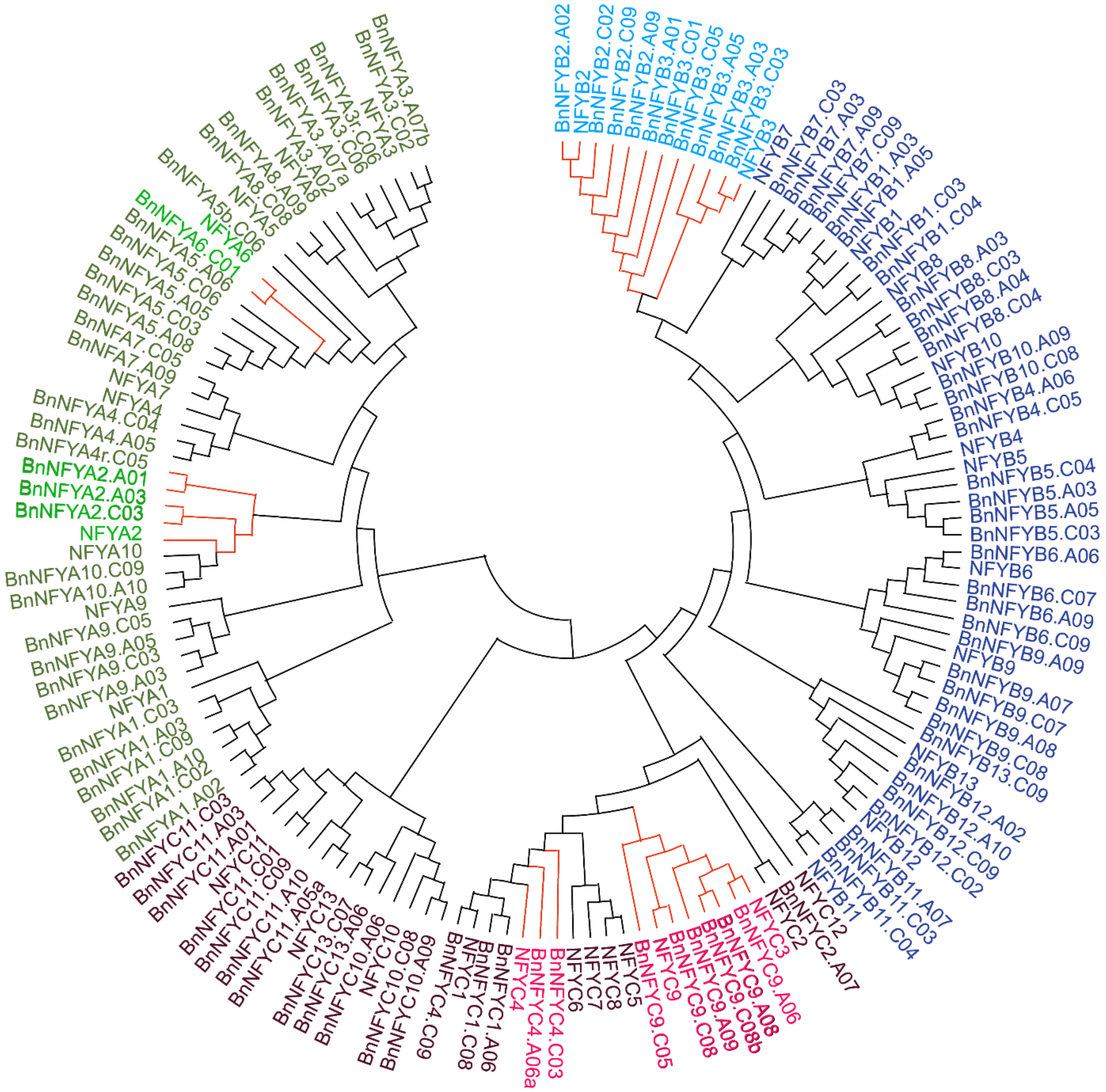


**Supplemental Figure 7. Phylogenetic tree constructed from the alignment of NF-Y homologs in *A. thaliana* and *B. napus*.** NF-YA, NF-YB and NF-YC homologs are coloured in green, blue and red, respectively. Branches marked in red indicate *A. thaliana* NF-Y proteins involved in *FT* regulation and their orthologs in *B. napus*.

| Query ID | Subject ID | Identity | Alignment Length | Mismatches | Gap Opening | Query Start | Query End | Subject Start | Subject End | E-value | Bit Score |
|---|---|---|---|---|---|---|---|---|---|---|---|
| AT1G65480.1 | BnaA02T0156900ZS | **87,69** | **528** | 65 | 0 | 1 | 528 | 1 | 528 | 5,30E-191 | 631 |
| AT1G65480.1 | BnaC02T0200600ZS | **87,5** | **528** | 66 | 0 | 1 | 528 | 1 | 528 | 7,70E-190 | 628 |
| AT1G65480.1 | BnaA07T0282700ZS | **86,74** | **528** | 70 | 0 | 1 | 528 | 1 | 528 | 3,30E-185 | 615 |
| AT1G65480.1 | BnaC06T0323800ZS | **86,55** | **528** | 71 | 0 | 1 | 528 | 1 | 528 | 4,70E-184 | 612 |
| AT1G65480.1 | BnaA07T0365100ZS | **82,92** | **521** | 89 | 0 | 1 | 521 | 1 | 521 | 1,20E-159 | 544 |
| AT1G65480.1 | BnaC04T0181400ZS | **81,23** | **522** | 98 | 0 | 1 | 522 | 1 | 522 | 3,00E-150 | 517 |
| AT1G65480.1 | BnaC02T0302200ZS | **81,61** | 310 | 57 | 0 | 213 | 522 | 294 | 603 | 5,00E-84 | 311 |
| AT1G65480.1 | BnaC06T0428800ZS | **81,48** | 270 | 49 | 1 | 1 | 269 | 1 | 270 | 3,90E-68 | 259 |
| AT1G65480.1 | BnaC02T0302200ZS | 77,61 | 201 | 45 | 0 | 1 | 201 | 1 | 201 | 3,40E-43 | 177 |
| AT1G65480.1 | BnaA06T0273500ZS | 63,36 | 393 | 141 | 1 | 15 | 404 | 12 | 404 | 5,70E-37 | 157 |
| AT1G65480.1 | BnaC03T0559000ZS | 62,85 | 393 | 143 | 1 | 15 | 404 | 12 | 404 | 4,70E-35 | 150 |
| AT1G65480.1 | BnaA03T0012400ZS | 60,08 | 491 | 193 | 1 | 18 | 505 | 27 | 517 | 4,30E-34 | 147 |
| AT1G65480.1 | BnaC03T0016500ZS | 59,47 | 491 | 196 | 1 | 18 | 505 | 18 | 508 | 3,20E-31 | 138 |

**Supplemental Table 1.** Last results of FT against proteins predicted in B. napus ZS11. Identities in bold were over the threshold to consider FT like proteins. Alignment Length in bold distinguish FT like proteins from fragments encoded by pseudogenes.

| Gene | Alias | Synteny Group | Genome | family | species |
|---|---|---|---|---|---|
| AT5G12840 | NFYA1 | A | At | NFYA | *A.thaliana* |
| BnaA02G0045100ZS | BnaNFYA1.A02 | A | A | NFYA | *B.napus* |
| BnaA03G0050200ZS | BnaNFYA1.A03 | A | A | NFYA | *B.napus* |
| BnaA10G0226800ZS | BnaNFYA1.A10 | A | A | NFYA | *B.napus* |
| BnaC02G0051800ZS | BnaNFYA1.C02 | A | C | NFYA | *B.napus* |
| BnaC03G0058100ZS | BnaNFYA1.C03 | A | C | NFYA | *B.napus* |
| BnaC09G0532200ZS | BnaNFYA1.C09 | A | C | NFYA | *B.napus* |
| AT3G05690 | NFYA2 | B | At | NFYA | *A.thaliana* |
| BnaA01G0396200ZS | BnaNFYA2.A01 | B | A | NFYA | *B.napus* |
| BnaA03G0300100ZS | BnaNFYA2.A03 | B | A | NFYA | *B.napus* |
| BnaC03G0359700ZS | BnaNFYA2.C03 | B | C | NFYA | *B.napus* |
| BnaC05G0533300ZS | BnaNFYA2.C05 | B | C | NFYA | *B.napus* |
| AT1G72830 | NFYA3 | C | At | NFYA | *A.thaliana* |
| BnaA02G0203300ZS | BnaNFYA3.A02 | C | A | NFYA | *B.napus* |
| BnaA07G0257900ZS | BnaNFYA3.A07b | C | A | NFYA | *B.napus* |
| BnaA07G0335600ZS | BnaNFYA3.A07a | C | A | NFYA | *B.napus* |
| BnaC02G0270200ZS | BnaNFYA3.C02 | C | C | NFYA | *B.napus* |
| BnaC06G0394200ZS | BnaNFYA3.C06 | C | C | NFYA | *B.napus* |
| BnaC06G0394000ZS | BnaNFYA3d.C06 | C | C | NFYA | *B.napus* |
| AT2G34720 | NFYA4 | D | At | NFYA | *A.thaliana* |
| BnaA05G0097800ZS | BnaNFYA4.A05 | D | A | NFYA | *B.napus* |
| BnaC04G0121000ZS | BnaNFYA4.C04 | D | C | NFYA | *B.napus* |
| AT1G54160 | NFYA5 | E | At | NFYA | *A.thaliana* |
| BnaA05G0156300ZS | BnaNFYA5.A05 | E | A | NFYA | *B.napus* |
| BnaA06G0007400ZS | BnaNFYA5.A06 | E | A | NFYA | *B.napus* |
| BnaA08G0010100ZS | BnaNFYA5.A08 | E | A | NFYA | *B.napus* |
| BnaC03G0804900ZS | BnaNFYA5.C03 | E | C | NFYA | *B.napus* |
| BnaC06G0084400ZS | BnaNFYA5b.C06 | E | C | NFYA | *B.napus* |
| BnaC06G0136700ZS | BnaNFYA5.C06 | E | C | NFYA | *B.napus* |
| AT3G14020 | NFYA6 | F | At | NFYA | *A.thaliana* |
| BnaC01G0442400ZS | BnaNFYA6.C01 | F | C | NFYA | *B.napus* |
| AT1G30500 | NFYA7 | G | At | NFYA | *A.thaliana* |
| BnaA09G0409000ZS | BnaNFA7.A09 | G | A | NFYA | *B.napus* |
| BnaC05G0264300ZS | BnaNFA7.C05 | G | C | NFYA | *B.napus* |
| AT1G17590 | NFYA8 | H | At | NFYA | *A.thaliana* |
| BnaA09G0616800ZS | BnaNFYA8.A09 | H | A | NFYA | *B.napus* |
| BnaC08G0472300ZS | BnaNFYA8.C08 | H | C | NFYA | *B.napus* |
| AT3G20910 | NFYA9 | I | At | NFYA | *A.thaliana* |
| BnaA03G0365100ZS | BnaNFYA9.A03 | I | A | NFYA | *B.napus* |
| BnaA05G0347700ZS | BnaNFYA9.A05 | I | A | NFYA | *B.napus* |
| BnaC03G0445600ZS | BnaNFYA9.C03 | I | C | NFYA | *B.napus* |
| BnaC05G0377800ZS | BnaNFYA9.C05 | I | C | NFYA | *B.napus* |
| AT5G06510 | NFYA10 | J | At | NFYA | *A.thaliana* |
| BnaA10G0267400ZS | BnaNFYA10.A10 | J | A | NFYA | *B.napus* |

| | | | | | |
|---|---|---|---|---|---|
| BnaC09G0583300ZS | BnaNFYA10.C09 | J | C | NFYA | *B.napus* |
| AT2G38880 | NFYB1 | A | At | NFYB | *A.thaliana* |
| BnaA03G0188600ZS | BnaNFYB1.A03 | A | A | NFYB | *B.napus* |
| BnaA05G0068900ZS | BnaNFYB1.A05 | A | A | NFYB | *B.napus* |
| BnaC03G0221700ZS | BnaNFYB1.C03 | A | C | NFYB | *B.napus* |
| BnaC04G0077900ZS | BnaNFYB1.C04 | A | C | NFYB | *B.napus* |
| AT5G47640 | NFYB2 | B | At | NFYB | *A.thaliana* |
| AT5G47670 | NFYB6 | B | At | NFYB | *A.thaliana* |
| BnaA02G0309400ZS | BnaNFYB2.A02 | B | A | NFYB | *B.napus* |
| BnaA09G0215400ZS | BnaNFYB2.A09 | B | A | NFYB | *B.napus* |
| BnaA09G0215600ZS | BnaNFYB6.A09 | B | A | NFYB | *B.napus* |
| BnaC02G0419600ZS | BnaNFYB2.C02 | B | C | NFYB | *B.napus* |
| BnaC07G0261700ZS | BnaNFYB6.C07 | B | C | NFYB | *B.napus* |
| BnaC09G0250000ZS | BnaNFYB2.C09 | B | C | NFYB | *B.napus* |
| BnaC09G0250300ZS | BnaNFYB6.C09 | B | C | NFYB | *B.napus* |
| BnaA06G0414700ZS | BnaNFYB6.A06 | B | A | NFYB | *B.napus* |
| AT4G14540 | NFYB3 | C | At | NFYB | *A.thaliana* |
| BnaA01G0344800ZS | BnaNFYB3.A01 | C | At | NFYB | *B.napus* |
| BnaA03G0343900ZS | BnaNFYB3.A03 | C | A | NFYB | *B.napus* |
| BnaA05G0394900ZS | BnaNFYB3.A05 | C | A | NFYB | *B.napus* |
| BnaC01G0426200ZS | BnaNFYB3.C01 | C | C | NFYB | *B.napus* |
| BnaC03G0414900ZS | BnaNFYB3.C03 | C | C | NFYB | *B.napus* |
| BnaC05G0441600ZS | BnaNFYB3.C05 | C | C | NFYB | *B.napus* |
| AT1G09030 | NFYB4 | D | At | NFYB | *A.thaliana* |
| BnaA06G0054400ZS | BnaNFYB4.A06 | D | A | NFYB | *B.napus* |
| BnaC05G0067400ZS | BnaNFYB4.C05 | D | C | NFYB | *B.napus* |
| AT2G47810 | NFYB5 | E | At | NFYB | *A.thaliana* |
| BnaA03G0228800ZS | BnaNFYB5.A03 | E | A | NFYB | *B.napus* |
| BnaA05G0001800ZS | BnaNFYB5.A05 | E | A | NFYB | *B.napus* |
| BnaC03G0269200ZS | BnaNFYB5.C03 | E | C | NFYB | *B.napus* |
| BnaC04G0002000ZS | BnaNFYB5.C04 | E | C | NFYB | *B.napus* |
| AT2G13570 | NFYB7 | F | At | NFYB | *A.thaliana* |
| BnaA03G0391200ZS | BnaNFYB7.A03 | F | A | NFYB | *B.napus* |
| BnaA09G0104400ZS | BnaNFYB7.A09 | F | A | NFYB | *B.napus* |
| BnaC03G0484300ZS | BnaNFYB7.C03 | F | C | NFYB | *B.napus* |
| BnaC09G0105300ZS | BnaNFYB7.C09 | F | C | NFYB | *B.napus* |
| AT2G37060 | NFYB8 | G | At | NFYB | *A.thaliana* |
| BnaA03G0177300ZS | BnaNFYB8.A03 | G | A | NFYB | *B.napus* |
| BnaA04G0237700ZS | BnaNFYB8.A04 | G | A | NFYB | *B.napus* |
| BnaC03G0207500ZS | BnaNFYB8.C03 | G | C | NFYB | *B.napus* |
| BnaC04G0553200ZS | BnaNFYB8.C04 | G | C | NFYB | *B.napus* |
| AT1G21970 | NFYB9 | H | At | NFYB | *A.thaliana* |
| BnaA07G0126300ZS | BnaNFYB9.A07 | H | A | NFYB | *B.napus* |
| BnaA08G0241300ZS | BnaNFYB9.A08 | H | A | NFYB | *B.napus* |
| BnaA09G0465400ZS | BnaNFYB9.A09 | H | A | NFYB | *B.napus* |

| | | | | | |
|---|---|---|---|---|---|
| BnaC07G0185300ZS | BnaNFYB9.C07 | H | C | NFYB | *B.napus* |
| BnaC08G0270100ZS | BnaNFYB9.C08 | H | C | NFYB | *B.napus* |
| AT3G53340 | NFYB10 | I | At | NFYB | *A.thaliana* |
| BnaA09G0495300ZS | BnaNFYB10.A09 | I | A | NFYB | *B.napus* |
| BnaC08G0334000ZS | BnaNFYB10.C08 | I | C | NFYB | *B.napus* |
| AT2G27470 | NFYB11 | J | At | NFYB | *A.thaliana* |
| BnaA07G0155000ZS | BnaNFYB11.A07 | J | A | NFYB | *B.napus* |
| BnaC03G0276400ZS | BnaNFYB11.C03 | J | C | NFYB | *B.napus* |
| BnaC04G0206800ZS | BnaNFYB11.C04 | J | C | NFYB | *B.napus* |
| AT5G08190 | NFYB12 | K | At | NFYB | *A.thaliana* |
| BnaA02G0028200ZS | BnaNFYB12.A02 | K | A | NFYB | *B.napus* |
| BnaA10G0255300ZS | BnaNFYB12.A10 | K | A | NFYB | *B.napus* |
| BnaC02G0030900ZS | BnaNFYB12.C02 | K | C | NFYB | *B.napus* |
| BnaC09G0569100ZS | BnaNFYB12.C09 | K | C | NFYB | *B.napus* |
| AT5G23090 | NFYB13 | L | At | NFYB | *A.thaliana* |
| BnaC09G0063500ZS | BnaNFYB13.C09 | L | C | NFYB | *B.napus* |
| AT3G48590 | NFYC1 | A | At | NFYC | *A.thaliana* |
| BnaA06G0165700ZS | BnaNFYC1.A06 | A | A | NFYC | *B.napus* |
| BnaC08G0290400ZS | BnaNFYC1.C08 | A | C | NFYC | *B.napus* |
| AT1G56170 | NFYC2 | B | At | NFYC | *A.thaliana* |
| BnaA07G0203700ZS | BnaNFYC2.A07 | B2 | A | NFYC | *B.napus* |
| AT1G54830 | NFYC3 | C | At | NFYC | *A.thaliana* |
| AT5G63470 | NFYC4 | D | At | NFYC | *A.thaliana* |
| BnaA06G0286000ZS | BnaNFYC4.A06a | D | A | NFYC | *B.napus* |
| BnaC03G0542200ZS | BnaNFYC4.C03 | D | C | NFYC | *B.napus* |
| BnaC09G0077900ZS | BnaNFYC4.C09 | D | C | NFYC | *B.napus* |
| AT5G50470 | NFYC7 | E | At | NFYC | *A.thaliana* |
| AT5G50480 | NFYC6 | E | At | NFYC | *A.thaliana* |
| AT5G50490 | NFYC5 | E | At | NFYC | *A.thaliana* |
| AT5G27910 | NFYC8 | F | At | NFYC | *A.thaliana* |
| AT1G08970 | NFYC9 | G | At | NFYC | *A.thaliana* |
| BnaA06G0054000ZS | BnaNFYC9.A06 | G | A | NFYC | *B.napus* |
| BnaA08G0297000ZS | BnaNFYC9.A08 | G | A | NFYC | *B.napus* |
| BnaA09G0663800ZS | BnaNFYC9.A09 | G | A | NFYC | *B.napus* |
| BnaC05G0066900ZS | BnaNFYC9.C05 | G | C | NFYC | *B.napus* |
| BnaC08G0189800ZS | BnaNFYC9.C08b | G | C | NFYC | *B.napus* |
| BnaC08G0528300ZS | BnaNFYC9.C08 | G | C | NFYC | *B.napus* |
| AT1G07980 | NFYC10 | H | At | NFYC | *A.thaliana* |
| BnaA06G0046600ZS | BnaNFYC10.A06 | H | A | NFYC | *B.napus* |
| BnaA09G0668200ZS | BnaNFYC10.A09 | H | A | NFYC | *B.napus* |
| BnaC08G0533200ZS | BnaNFYC10.C08 | H | C | NFYC | *B.napus* |
| AT3G12480 | NFYC11 | I | At | NFYC | *A.thaliana* |
| BnaA01G0367300ZS | BnaNFYC11.A01 | I | A | NFYC | *B.napus* |
| BnaA03G0328100ZS | BnaNFYC11.A03 | I | A | NFYC | *B.napus* |
| BnaA05G0425500ZS | BnaNFYC11.A05a | I | A | NFYC | *B.napus* |

| | | | | | |
|---|---|---|---|---|---|
| BnaA05Z0425400ZS | BnaNFYC11.A05b | I | A | NFYC | *B.napus* |
| BnaC01G0458700ZS | BnaNFYC11.C01 | I | C | NFYC | *B.napus* |
| BnaC03G0393600ZS | BnaNFYC11.C03 | I | C | NFYC | *B.napus* |
| BnaA10G0258900ZS | BnaNFYC11.A10 | new | A | NFYC | *B.napus* |
| BnaC09G0573000ZS | BnaNFYC11.C09 | new | C | NFYC | *B.napus* |
| AT5G38140 | NFYC12 | J | At | NFYC | *A.thaliana* |
| AT5G43250 | NFYC13 | K | At | NFYC | *A.thaliana* |
| BnaA06G0442500ZS | BnaNFYC13.A06 | K | A | NFYC | *B.napus* |
| BnaC07G0218200ZS | BnaNFYC13.C07 | K | C | NFYC | *B.napus* |

**Supplemental Table 2.** NF-Y genes in Brassica napus ZS11 and A. thaliana. Syntenic genes per group are annotated to the same letter.

| Application | Primer name | Sequence |
|---|---|---|
| qPCR in *B. napus* | BnaFT.A2-qF | GGTATTCATCGTATCGTGCTCG |
| | BnaFT.A2-qR | CAAGTTATTAAAAGAAGAAGAGGCTC |
| | BnaFT.C2-qF | GGTATTCATCGTATCGTGCTG |
| | BnaFT.C2-qR | GTTATTAAAAGAAGAAGAGGCTCATC |
| | BnaFT.A7-qF | CCACCTCGGGAATTCATCGTC |
| | BnaFT.A7-qR | CCATGACCCATCGATCTAAG |
| | BnaFT.C6-qF | CAAACGGTGTATGAACCAGG |
| | BnaFT.C6-qR | TCTAAGGAAGAAGCCCATCG |
| | BnaNFT.A7-qF | CGAGAGACCCTCTTATCGTAGG |
| | BnaNFT.A7-qR | AATCTCAACCGTTGGTTTGTTC |
| | BnaCFT.C4-qF | CGAGAGATCCTCTTGTGCTTGC |
| | BnaCFT.C4-qR | GATCTCGACCGTTGGTTTATTT |
| | BnaCO.A10-qF | ACGTATGGCTCCTCAGGAAGTCAC |
| | BnaCO.A10-qR | TCTGAATTAGAGGTTCAGGTAGTTTCT |
| | BnaCO.C09-qF | TAAACAAGACTGCATCGTACCAGAGA |
| | BnaCO.C09-qR | GTCAGTTTCCATTGATGGATTGTATG |
| | BnaENTH-qF | GTTTAGACCCGTTGCTGCTC |
| | BnaENTH-qR | TTGTCCATCTCAGCCATTTG |
| qPCR in *A.thaliana* | RT-BnFT_cDNA_F | GGTGGAGAAGACCTAAGGAA |
| | RT-BnFT_cDNA_R1 | GGTTCATACACTGTTTGCCT |
| | RT-BnCFT_cDNA_F | TGTCCCACTTCCGGAATTCA |
| | RT-BnCFT_cDNA_R | TCCTCCACAGCCATTCTCTC |
| | RT-BnNFT_cDNA_F | ACGAGAATCCATGTCCCACA |
| | RT-BnNFT_cDNA_R | TGAAGTAAACAGCAGCCACG |
| | RT-FT_cDNA_F | GGTGGAGAAGACCTCAGGAA |
| | RT-FT_cDNA_R | ACCCTGGTGCATACACTGTT |
| | BASTA_qPCR_F | CCAGTTCCCGTGCTTGAA |
| | BASTA_qPCR_R | AGAGCGTGGTCGCTGTCAT |
| | PP2A_qPCR_F | ACACAATTCGTTGCTGTCTTCT |
| | PP2A_qPCR_R | TGCTTGGTGGAGCTAAGTGA |
| Complementation with plasmid driven by FT Block (C+A) | BnaFT.A2C2-pFT-F | TGGTGATATCAAGCTTATGTCTTTAAGTAATAGAGATCCTCTTG |
| | BnaFT.A2-pFT-R | GATCGGGGAAATTCGAGCTCCTAACTTCTTCGTCCTCCG |
| | BnaFT.C2-pFT-R | GATCGGGGAAATTCGAGCTCCTAACTTCTTCGTCCTCCGCAGC |
| | BnaFT.A7C6-pFT-F | TGGTGATATCAAGCTTATGTCTGTAAATAACAGAGATCCTCT |
| | BnaFT.A7C6m-pFT-R | GATCGGGGAAATTCGAGCTCCTAAGTTCTTCGTCCTCCG |
| | BnaFT.C6C7m-pFT-R | GATCGGGGAAATTCGAGCTCCTAACATCTTCGTCCTCCG |
| | BnaNFT.A7-pFT-F/ | TGGTGATATCAAGCTTATGTCATTAAGTCCGAGAGACCCT |
| | BnaNFT.A7-pFT-R | GATCGGGGAAATTCGAGCTCCTACGAGGTCCTTCTTCCTCCG |
| | BnaCFT.C4-pFT-F | TGGTGATATCAAGCTTATGTCTTTAAGTCCGAGAGATCCTC |
| | BnaCFT.C4-pFT-R | GATCGGGGAAATTCGAGCTCCTATGTTCTTCTTCCTCCACAGCCA |
| | Plasmid-V-F1 | CACAGAGAAACCACCTGTTTGTT |
| | Plasmid-V-R1 | TATGATAATCATCGCAAGACCG |
| ComplemenComplementation with | BnaFT.A7C6-pFD-F | AGGTGGTGGAAGTTACCCTTACGATGTGCCTGATTACGCTGGAAGTTCTGTAAATAACAGAGATCCTCTTG |

| | | |
|---|---|---|
| plasmid driven by FD promoter | pFD-2HA-F | CTTCTGTTCTCTTTTCCAATGTACCCATACGATGTGCCTGATTACGCTGGAGGTGGTGGAAGTTACCCTTACG |
| | BnaFT.A7-pFD-R | CAAGGACTTGTAGATTTCCTAAGTTCTTCGTCCTCCG |
| | BnaFT.C6-pFD-R | CAAGGACTTGTAGATTTCCTAACATCTTCGTCCTCCG |
| | At.FT-pFD-F | AGGTGGTGGAAGTTACCCTTACGATGTGCCTGATTACGCTGGAAGTTCTATAAATATAAGAGACCCTCT |
| | At.FT-pFD-R | CAAGGACTTGTAGATTTCCTAAAGTCTTCTTCCTCCGCA |
| | Plasmid-V-F2 | ACCGGCTAAAGTCAAGAACCTCT |
| | Plasmid-V-R2 | CCGGGTCTTTTGTTTTACATCTTC |
| | 2HA-tag | TACCCATACGATGTGCCTGATTACGCTGGAGGTGGTGGAAGTTACCCTTACGATGTGCCTGATTACGCT |
| Tobacco infiltration, reporter vectors construction | pFT-5.7K-F | CGA ATT GGG TAC AGT ACT CCTCTCTTCGAATTACATTCGTATGA |
| | pFT-5.7K-R | TCG CGT TTC ACC ATG G CTTTGATCTTGAACAAACAGGTGGT |
| | pA2-12K-P1-F | CGAATTGGGTACAGTACTGCTATCAATAGTAATTCGATTTCTATGAGC |
| | pA2-12K-P1-R | ACCAGATGATGCCTGCGTCTATG |
| | pA2-12K-P2-F | GCAGGCATCATCTGGTGAGAAC |
| | pA2-12K-P2-R | TCGCGTTTCACCATGGCTTTGATCTAAAACAAACAGGTGG |
| | pA7-12K-P1-F | CGAATTGGGTACAGTACTGTGCTTTAAACTAGTGACCAGGAG |
| | pA7-12K-P1-R | TCCAAACTTCTTTGCAACAGACAAAGG |
| | pA7-12K-P2-F | GCAAAGAAGTTTGGATTCACTCAG |
| | pA7-12K-P2-R | TCGCGTTTCACCATGGCTCTGATCTAAAACAAACAGGTTG |
| | pC2-21K-P1-F | CGAATTGGGTACAGTACTGCACAAAAGTTACGTTTTGTTACAGC |
| | pC2-21K-P1-R | CAGAAGACTTCTCCCAACAGAC |
| | pC2-21K-P2-F | GTTGGGAGAAGTCTTCTGTGTTAG |
| | pC2-21K-P2-R | TAGACTATGCTGCCCTTAATCTCTTCG |
| | pC2-21K-P3-F | GGGCAGCATAGTCTAGTTTTGAG |
| | pC2-21K-P3-R | GTTAATCTTGTGTTTTCTTGTAGTGAATC |
| | pC2-21K-P4-F | ACAAGAAAACACAAGATTAACGACG |
| | pC2-21K-P4-R | TCGCGTTTCACCATGGCTTTGATCTAAAACAAACAGGTGG |
| | pC6-1.8K-F | CGA ATT GGG TAC AGT ACT GAGCCATTAGATTCGTATGATCACG |
| | pC6-1.8K-R | TCG CGT TTC ACC ATG G CTCTGATCTAAAACAAACAGGTTGTTTC |
| Tobacco infiltration, effector vectors construction | adScaI-35S-F | CGAATTGGGTACAGTACT GTGGAGCACGACACAC |
| | adSacI-CO-R | GATCGGGGAAATTCGAGCTC TCAGAATGAAGGAACAATCCC |
| | adSacI-BnCO.A10/C9-R | GATCGGGGAAATTCGAGCTC TTATTTTTGGCCATAGAATGAAGG |
| Tobacco infiltration, plasmid verification | V-629-F1 | TGAAGCAACTCCTCGAAAAAGC |
| | V-629-R1 | CAATTCCACACAACATACGAGCC |
| | V-629-F2 | GTGCTGCAAGGCGATTAAGTTG |
| | V-629-R2 | ACGAGTGCTTGAGGGAGGTGAC |

**Supplemental Table 3. Oligonucleotide sequences used in this study**

| Arabidopsis (TAIR) | | Brassica napus | | | Schrenkiella parvula | | Brassica rapa | | Brassica oleracea | |
|---|---|---|---|---|---|---|---|---|---|---|
| Name | Araport 11 | Name | Gene ID ZS11 v0.0 | Gene ID Darmor | Name | Gene ID v2.2 | Name | Gene ID BraZ1 | Name | Gene ID 2J.m1 |
| FT | AT1G65480 | BnaFT.A2 | BnaA02G0156900ZS | BnaA02g12130D | | | BrFT.A2 | A02p18180 | | |
| | | BnaFT.C2 | BnaC02G0200600ZS | BnaC02g45250D | | | | | BoFT.C2 | BolC02g021840 |
| | | BnaFT.A7 | BnaA07G0282700ZS | BnaA07g25310D | | | BrFT.A7 | A07p35130 | | |
| | | BnaFT.C6 | BnaC06G0323800ZS | BnaC06g27090D | | | | | BoFT.C6 | BolC06g036950 |
| NFT | absent | BnaNFT.A7 | BnaA07G0365100ZS | BnaA07g33120D | SpNFT | Sp5g32040 | BrNFT.A7 | A07p44520 | | |
| | | BnaNFT.C6 | BnaC06T0428800ZS | not predicted | | | | | BoNFT.C6 | BolC06g048270 |
| | | BnaNFT.A2 | absent | absent | | | BrNFT.A2 | A02p26660 | | |
| | | BnaNFT.C2 | BnaC02T0302200ZS | BnaC02g23820D | | | | | BoNFT.C2 | BolC02g033320 |
| CFT | absent | BnaCFT.C4 | BnaC04G0181400ZS | BnaC04g14850D | | absent | | absent | BoCFT.C4 | BolC04g020210 |
| TSF | AT4G20370 | absent | absent | absent | SpTSF | Sp7g18730 | | absent | | absent |

**Supplemental Table 4.** Summary Table of *FT* like genes in *A. thaliana*, *B. napus* var. ZS11 and Darmor, *S. parvul*a, *B. rapa* and *B.oleracea.* Pseudogenes are marked in red.
